# The tyrosine kinase inhibitor GNF-7 targets senescent cells through allosteric activation of GCN2

**DOI:** 10.1101/2025.08.29.673062

**Authors:** Gema Lopez-Pernas, Matilde Murga, Wareed Ahmed, Maria Haggblad, Elena Jiménez-Ortega, Alicia G. Serrano, Carlota Cardona, Mario Lopez-Prieto, Samuele Fisicaro, Jorge Mota-Pino, Belén Navarro-Gonzalez, Eduardo Zarzuela, Marta E. Anton, Louise Lidemalm, Sonia Martinez, Marta Isasa, Joaquín Pastor, Rafael Fernandez-Leiro, Daniela Huhn, Oscar Fernández-Capetillo

## Abstract

The one-two-punch approach refers to the sequential administration of two different chemotherapies, the second of which targets cancer cells that resisted the initial treatment. To find such a second punch, we performed a chemical screen to find drugs that are preferentially toxic for cells with an activated DNA damage response (DDR). This screen identified the tyrosine kinase inhibitor GNF-7 as a top hit. Subsequent work revealed that GNF-7 is a potent senolytic, even when senescence is triggered by therapies that do not activate the DDR. Consistently, GNF-7 is highly efficacious to kill cancer cells previously treated with CDK4/6 inhibitors, including in patient-derived organoids and mouse xenografts. Surprisingly, the senolytic effect of GNF-7 is not mediated by the inhibition of a tyrosine kinase (TK), but rather by the activation of GCN2, an effect previously reported for other TK inhibitors. Together, our study reports the discovery of a novel senolytic agent that strongly synergizes with CDK4/6 inhibitors when applied sequentially and expands our understanding of the mechanisms behind the anticancer effects of TK inhibitors.

## Introduction

Cancer chemotherapies often fail to kill all malignant cells, which drives tumor relapse months or years after the initial treatment, leading to secondary disease of worse prognosis. In this context, there is significant effort to find therapies that target therapy-resistant cancer cells, which are often in growth-arrested cell states such as senescence^1,2^ or drug-tolerant persistence (DTP)^3,4^. In particular, the discovery of drugs that can target therapy-induced senescent cells (TIS), also known as senolytics, has gained significant attention, as many of the therapies used in oncological practice induce senescence^5,6^, and also since senolytic drugs could have beneficial effects on other age-related pathologies besides cancer^1,2,7^. Despite being unable to grow, senescent cells are metabolically active and can fuel secondary tumor growth through the pro-inflammatory and immunosuppressive senescence associated secretory phenotype (SASP)^8,9^. In addition, senescent cells are resistant to the induction of apoptosis, which is due to the increased activity of pro-survival signaling pathways as well as the expression of anti-apoptotic factors from the BCL-2 family of proteins^10^.

Senolytic therapies have tried to exploit intrinsic features of senescent cells. For instance, early works targeted BCL-2, BCL-XL and BCL-W with selective inhibitors like navitoclax (ABT263) ^11^, or its second generation venetoclax (ABT-199), which presents reduced side effects on platelet counts^12^. However, not all senescent cells upregulate BCL-2 family proteins and are thus insensitive to these agents^13^. Other senolytic strategies include peptides targeting the p53/FOXO4 interaction^14^, cardiac glycosides^15,16^ and CAR-T cells directed against senescent membrane receptors^17^. Besides these targeted therapies, the term “senolytics” was first described in a manuscript describing the effects of the multitarget tyrosine kinase inhibitor dasatinib, which are potentiated by the flavonoid quercetin^18^. While dasatinib is a frequently used senolytic agent, the mechanism of this phenomenon remains unknown.

In what regards to the use of senolytics in cancer therapy, the emergent view is that these agents would be best suited in a “one-two-punch” regimen, whereby a first pro-senescent treatment is followed by a senolytic drug to maximize the clearance of cancer cells^19^. A recent example of this approach is the use of lysomotropic agents to target cells previously treated with CDK4 inhibitors, which exploits the increased reliance of senescent cells on lysosomal activity^20^. Here, we aimed to discover novel agents that are preferentially toxic for cancer cells previously treated with chemotherapy, in order to maximize the efficacy of the treatment.

## Results

### A chemical screen targeting DDR+ cells identifies senolytic compounds

Our initial aim was to identify drugs that preferentially kill cells with an activated DNA damage response (DDR), which is a hallmark of cancer cells treated with genotoxic chemotherapies. To do so, we used a previously described system whereby a fragment of TOPBP1 that allosterically activates the ATR kinase is fused to a sequence from the Estrogen Receptor (ER) (TopBP1^ER^). In cells expressing this fragment, the DDR can be activated by addition of 4-hydroxytamoxifen (4-OHT)^21^. We generated clones of MCF7 human breast cancer cells expressing TOPBP1^ER^ (MCF7^ER^), as well as EGFP or mCherry to enable their tracking. The screening was performed in 384-well plates, where we seeded MCF7^ER/EGFP^ cells previously treated with 4-OHT for 16h (DDR^ON^), together with MCF7^mCherry^ cells that had not been exposed to 4-OHT (DDR^OFF^) at a 1:2.5 ratio. 16h after seeding, co-cultures were exposed to 10 μM of 5,285 drugs from the chemical library for 40 additional h, after which the proportion of DDR^ON^ and DDR^OFF^ cells was analyzed by High Throughput Microscopy (**Fig. 1A, B** and **Table S1**). The library of choice was the one defined by the Drug Repurposing Hub initiative^22^, which contains medically approved drugs, chemicals that have gone through early phase clinical trials and tool compounds.

**Figure 1.**
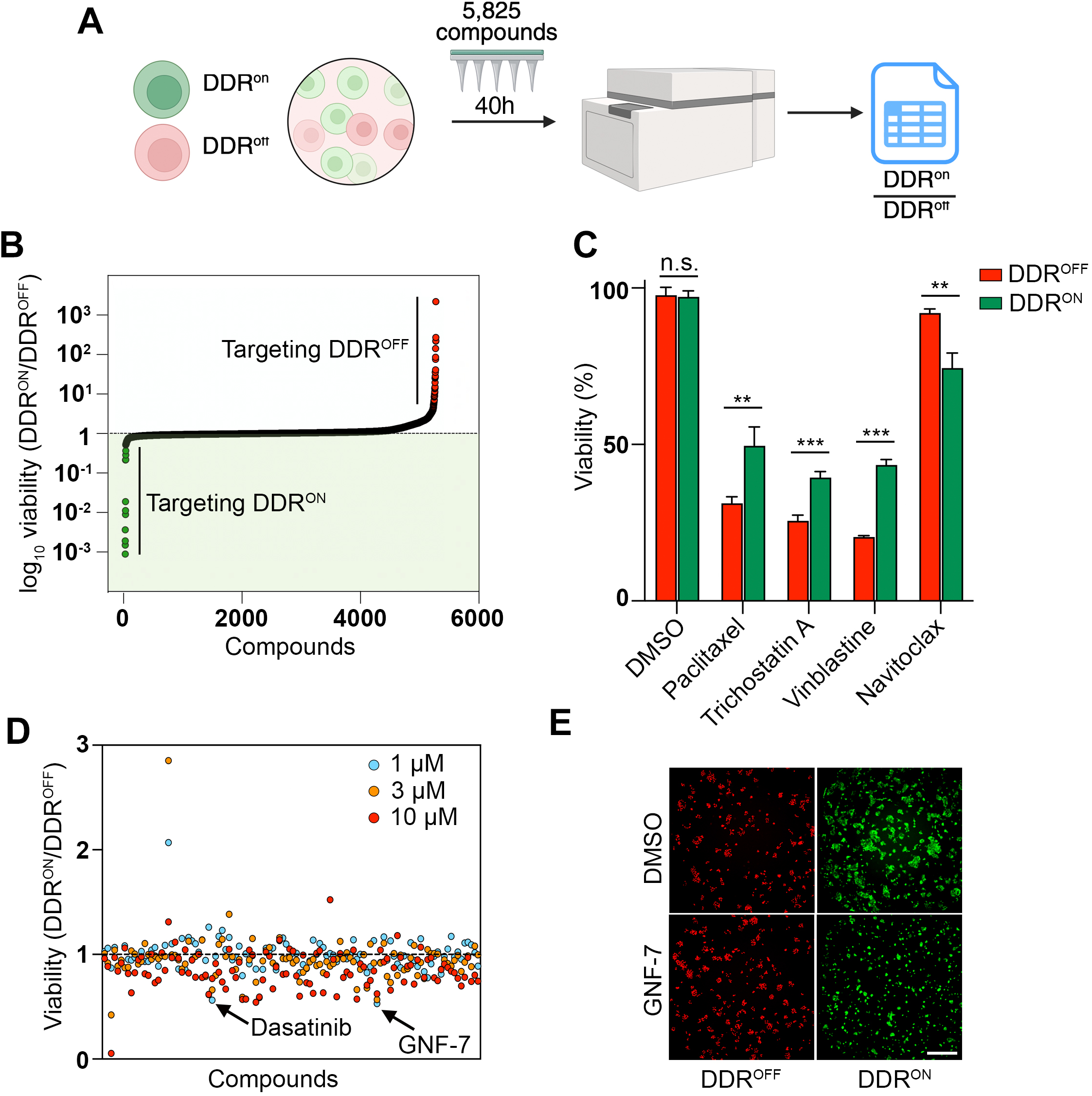
A chemical screen identifies GNF-7 as a senolytic drug. (**A**) Schematic pipeline of the chemical screen. Control (mCherry, DDR^OFF^) and TopBP1^ER^ expressing (GFP, DDR^ON^) MCF7 cells were treated with 50nM 4-OHT for 16 h and mixed at a 1:2.5 ratio. 6 h after, cells were exposed to exposed to 5,285 drugs from the chemical library (10 μM, 40 h treatment, triplicates). Viable DDR^ON^ and DDR^OFF^ cells were quantified using HTM. (**B**) Distribution of screen data. Hits highlighted in red are those preferentially targeting DDR^OFF^ cells, while the ones in green preferentially target DDR^ON^ cells. **(C)** Normalized viability, as evaluated by HTM-mediated quantification of nuclei, of MCF-7 DDR^OFF^ (red) and DDR^ON^ (green) treated with the indicated compounds for 24 h. (**D**) Results from the dose-response validation screen, illustrating the impact of the primary hits in DDR^OFF^/DDR^ON^ ratios at the indicated doses. The experiment used the same conditions as defined in (A). (**E**) Representative immunofluorescence of the data shown (D) after treatment with 1 μM GNF-7 for 72 h or DMSO as a control. Scale bar (white) indicates 100 μm. n.s.: non-significant, \*\**P* < 0.01; \*\*\**P* < 0.001; *t*-test.

Most compounds did not affect viability in either cell population. As for toxic chemicals (defined as the 10% percentile of compounds with a bigger impact on cell viability), these were overall more toxic for DDR^OFF^ cells (mean viability 38,57%) than for DDR^ON^ cells (mean viability 57,19%). Given that exposure to 4-OHT triggers senescence in MCF7^ER^ cells^21^, we hypothesized that this observation could reflect an increased resistance to apoptosis of the senescent cells. Supporting this view, co-culture experiments of DDR^ON^/DDR^OFF^ cells revealed that whereas DDR^ON^ cells were indeed resistant to chemotherapies such as paclitaxel, vinblastine or trichostatin A, they were sensitive to the senolytic navitoclax (**Fig. 1C**). These results suggested that our screen could be a valid platform for the discovery of novel senolytics, by focusing on drugs that preferentially killed DDR^ON^ cells.

Hits were selected from the primary screen as those affecting the viability of DDR^ON^ cells more than 20% and decreasing the DDR^ON^/DDR^OFF^ ratio below 0.8. In addition, we removed compounds that affected the analysis due to their intrinsic fluorescence such as doxorubicin. On this basis, we selected 112 compounds for validation in a dose-response screen conducted at 1, 3 and 10μM, following the same pipeline defined above. The results from this secondary screen (**Fig. 1D** and **Table S2**) confirmed the preferential toxicity of senolytic compounds such as dasatinib for DDR^ON^ cells. The top hit from the validation screen was the tyrosine kinase inhibitor GNF-7 (**Fig. 1E**), originally developed as an inhibitor of ABL1 for the treatment of chronic myeloid leukemias^23^. Given its potency and good pharmacological properties, we set to determine whether GNF-7 was a novel senolytic drug.

### GNF-7 is a broad spectrum senolytic drug

Next, we set to determine efficacy of GNF-7 as a senolytic in several independent models. First, we tested GNF-7 in an additional model of DDR-induced senescence. To do so, we induced senescence in MCF7 cells by exposure to doxorubicin for 5 days. At this time, a treatment with dasatinib or GNF-7 was preferentially toxic for senescent MCF7 cells (**Fig. 2A**). Next, we moved to HT-1080 human fibrosarcoma cells, which are a widely characterized model for the study of cellular senescence^14,24^. To induce senescence in a controlled manner, we generated a clone of HT-1080 cells harboring doxycycline (dox)-inducible expression of a dominant negative fragment of the telomeric protein TRF2 (TRF2Λ′N)^13^. In these cells (HT-1080^T^), a 6-day treatment with dox led to telomere-uncapping, activation of the DDR, senescence and sensitivity to navitoclax (**Fig. S1A, B**). Senescent HT-1080^T^ cells were similarly sensitive to GNF-7 regardless of whether senescence was induced by telomere uncapping or by the DNA damaging agent etoposide (**Fig. S1C**). A time-point analysis revealed that the senolytic effects of GNF-7 in HT-1080^T^ cells followed a similar kinetics as that of navitoclax (**Fig. S1D, E**).

**Figure 2.**
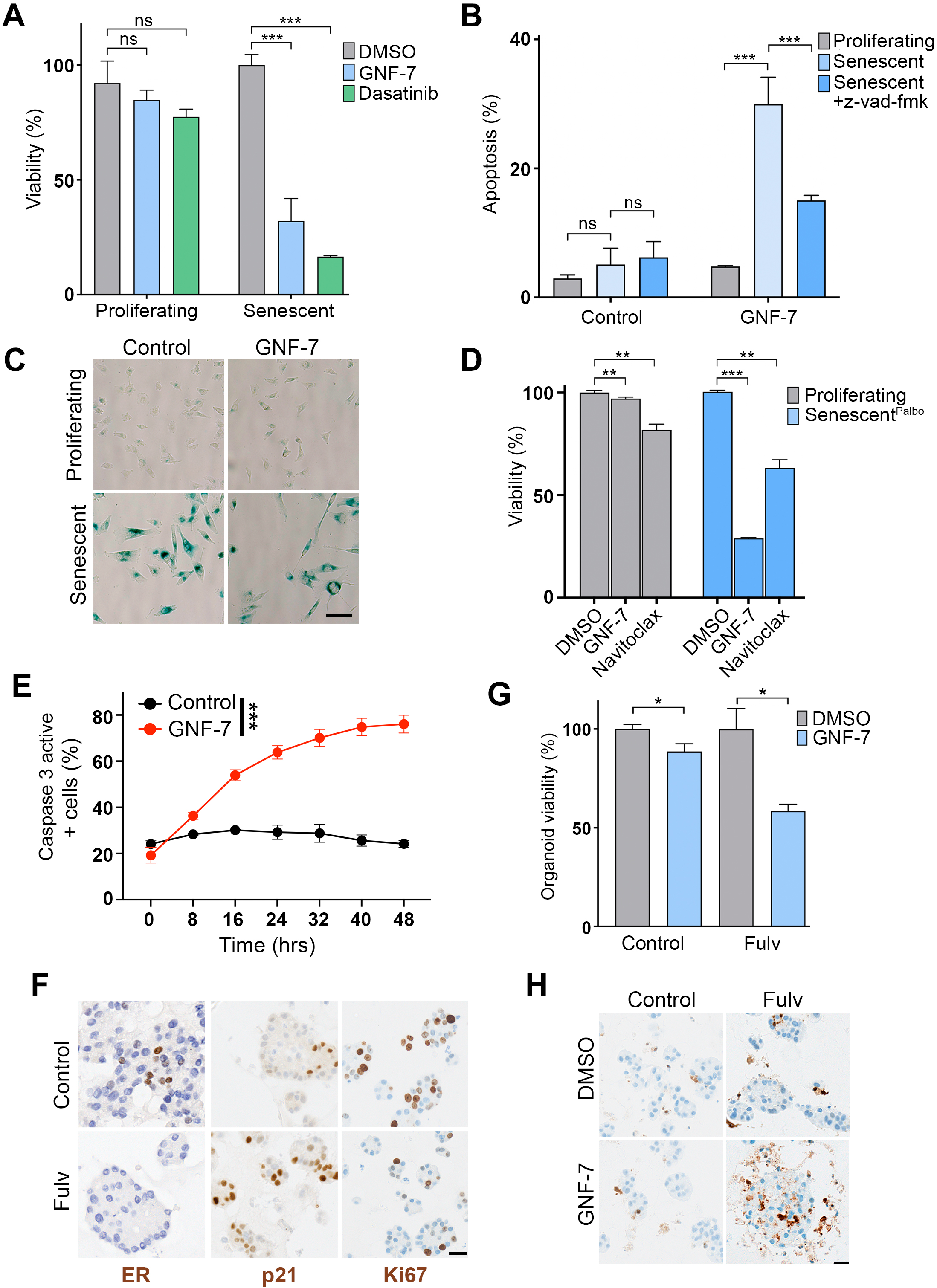
GNF-7 is a broad-spectrum senolytic agent. (**A**) Normalized cell viability (as quantified by HTM through counting nuclei) of proliferating and senescent MCF-7 cells after treatment with GNF-7 (200nM, blue) and dasatinib (10 μM, green) for 72 h. Senescence was induced by doxorubicin (50 nM) for 5 days. (**B**) Percentage of apoptotic cells (measured by FACS as Annexin V positive and PI negative) upon GNF-7 treatment in proliferating and senescent HT-1080 cells. Apoptosis was inhibited with the pan-caspase inhibitor Z-VAD-FMK (20 μM). Senescence was induced by palbociclib (5 μM) for 7 days. (**C**) Representative image of SA-β-Gal staining of proliferating and senescent HT-1080 cells upon treatment with 75 nM GNF-7 for 72 h. Scale bar (black) indicates 10 μm. Senescence was induced by palbociclib (5 μM) for 7 days. (**D**) Normalized viability, as evaluated by CellTiter-Glo, in proliferating or senescent HT-1080 cells, upon treatment with 200nM GNF-7 or 5 μM Navitoclax for 72 h. Senescence was induced by palbociclib (5 μM) for 7 days. (**E**) Percentage of caspase 3 active senescent HT-1080 cells treated with GNF-7 (200 nM) or DMSO as a control for 48 h. (**F**) Representative immunohistochemical images of ER, p21 and KI-67 expression in ER+ breast cancer organoids after 1μM fulvestrant treatment of for 7 days. (**G**) Normalized organoid viability, as evaluated by HTM-mediated quantification of nuclei, upon treatment of the organoids with fulvestrant alone, or fulvestrant followed by a treatment with 500nM GNF-7 for 20 days. (**H**) Representative immunohistochemical images of cleaved caspase-3 expression in organoids treated as in (G). Scales bar (black) indicate 20 μm. n.s.: non-significant, \**P* < 0.05; \*\**P* < 0.01; \*\*\**P* < 0.001; *t*-test.

While performing the previous experiments, we consistently observed that GNF-7 also decreased cell numbers in cultures of proliferating cells. However, this was due to a cytostatic effect of the compound in triggering G1 arrest, as evidenced by an increase in the percentage of HT-1080 cells with a G1 DNA content upon exposure to GNF-7, together with a reduction in the percentage of replicating cells that incorporate ethynyl deoxyuridine (EdU) **(Fig. S2A, B**). In addition, RNA sequencing (RNAseq) analyses of proliferating HT-1080 cells treated with GNF-7 revealed a generalized downregulation of pathways related to cell growth (**Fig. S2C**). In contrast, treatment with GNF-7 triggered apoptosis in palbociclib-induced senescent HT-1080 cells, as measured by annexin staining, in a manner that was rescued by the pan-caspase inhibitor z-vad-fmk (**Fig. 2B, C**). Hence, GNF-7 has cytostatic effects in proliferating cells, but selectively triggers caspase-dependent apoptosis in senescent cells. Noteworthy, the senolytic effects of GNF-7 were more pronounced than those of navitoclax in HT-1080 cells treated with the CDK4/6 inhibitor palbociclib (**Fig. 2D**).

Time-lapse videos with a probe detecting Caspase-3/7 activity confirmed that GNF-7 triggered apoptosis on HT-1080 cells, only if these were previously treated with palbociclib (**Fig. 2E** and **Videos S1, S2**). To further test the senolytic effects of GNF-7 in a DDR-independent model, we used patient-derived tumor organoids from estrogen receptor positive (ER+) breast cancer. As we previously reported, treatment of ER+ breast cancer cells with the selective estrogen receptor degrader fulvestrant triggers a senescent phenotype^25^. Consistently, fulvestrant-treated tumor organoids had a reduction in KI67 positive cells together with an increase in cells with p21 expression (**Fig. 2F**). Importantly, GNF-7 was preferentially toxic for fulvestrant-treated organoids, where the treatment induced apoptosis as evidenced by an increase in activated caspase 3 (**Fig. 2G, H)**. In summary, these experiments identify GNF-7 as a novel senolytic compound with activity in a wide range of senescent models.

### G1 arrest and PI3K/AKT signaling modulate the response to GNF-7

Next, we set to determine the signaling pathways that modulate the cellular response to GNF-7. To do so, we performed a genomewide CRISPR/Cas9 screen in mouse embryonic stem cells (mES) treated with GNF-7 for 10 days. The library of choice was the mouse Toronto knockout (mTKO) CRISPR library^26^, which targets 19,463 murine genes with 5 sgRNAs/gene. Of note, we chose mES as a cell model since the absence of proliferation limits the selection of sgRNAs in senescent cells and because we have successful experience in using this model for genetic screens related to drug responses^27–29^. Bioinformatic analysis of sgRNA fold changes at day 10 of GNF-7 treatment using MAGeCK^30^ revealed several interesting features.

Among the genes targeted by sgRNAs that were depleted in GNF-7-treated cells, there was an enrichment for those whose depletion leads to G1 arrest such as *Ccne1*, *Ccnc*, *Usp48*, *Ilk or Tln1* (**Fig. 3A, B**). Consistently, CRISPR-mediated depletion of ILK or TLN1 arrested HT-1080 cells in G1 and increased their sensitivity to GNF-7 (**Fig. S3A-C**). Conversely, CCNE1 overexpression promoted S-phase entry (as measured by EdU incorporation) and reduced GNF-7 toxicity in RPE cells (**Fig. S3 D-F**). Importantly, arresting HT-1080 cells in G1 through a double-thymidine block sensitized them to GNF-7 (**Fig. 3** **C, D**). Thus, in contrast to most chemotherapies that target rapidly growing cells, GNF-7 is preferentially toxic for growth arrested cells.

**Figure 3.**
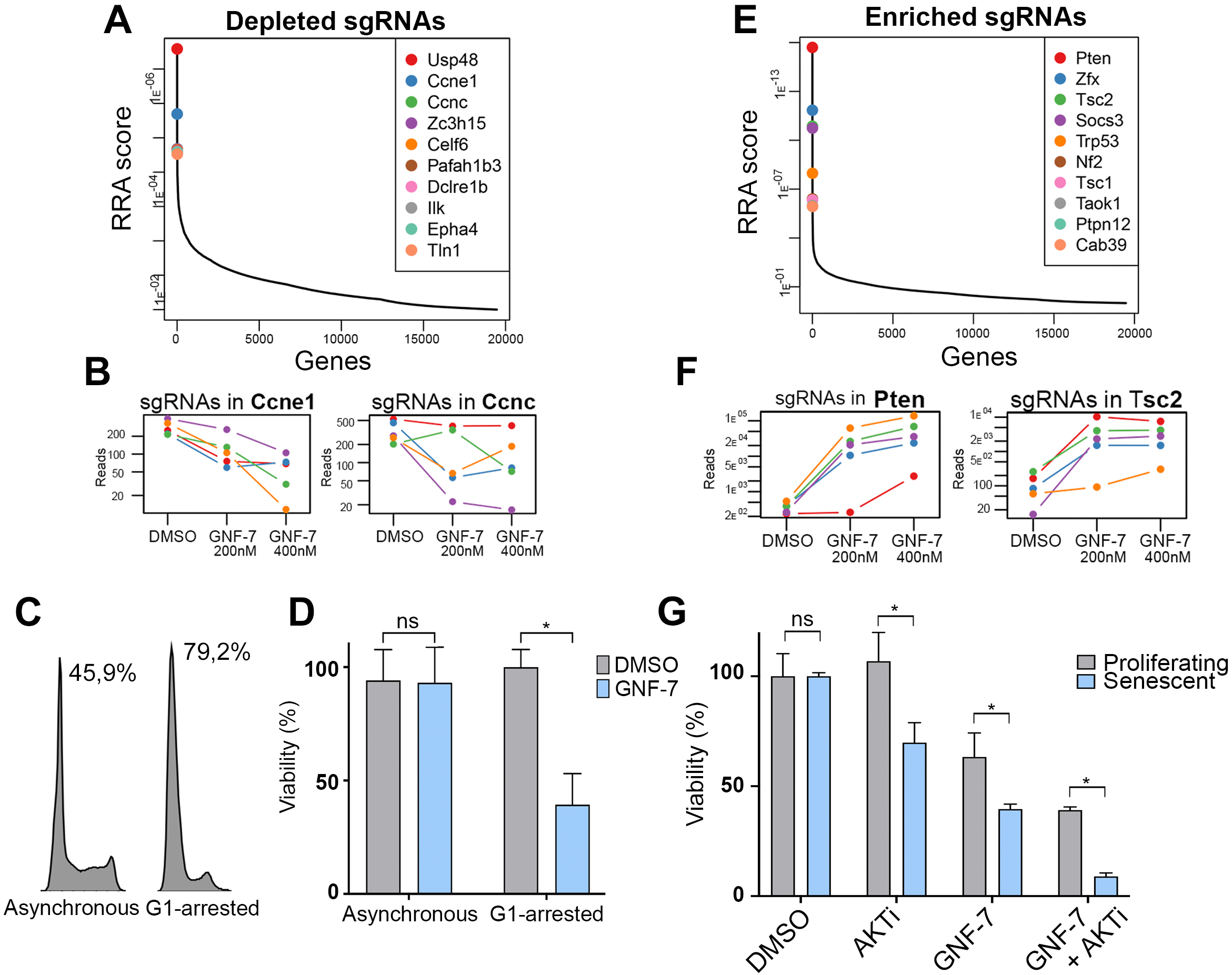
A CRISPR screen to identify the determinants of GNF-7 toxicity. (**A**) Robust Rank Aggregation scores (RRA score, left Y axis) of depleted sgRNAs (negatively selected), calculated using the MAGeCK computational tool. The plot depicts the top 10 genes with depleted sgRNAs in the GNF-7 (combining data from 200 and 400nM treatments) compared to the DMSO control. (**B**) Representative examples of the read counts of the sgRNAs targeting two of the top enriched hits. **(C)** Cell cycle distribution (based on DNA content) in proliferating or G1-arrested HT-1080 cells. Cells were arrested by a double-thymidine block, and the numbers indicate the percentage of cells in G1. **(D)** Normalized viability of the cells described in (C) upon treatment with 400 nM GNF-7 for 48 h, as measured by CellTiter-Glo. (**E**) RRA scores for the top 10 genes with enriched sgRNAs (positively selected). (**F**) Representative examples of the read counts of the sgRNAs targeting two of the top enriched hits. **(G)** Normalized viability, as evaluated with CellTiter-Glo, of proliferating and senescent HT-1080 cells upon treatment with 100 nM GNF-7, alone or in combination with 2 μM Capivasertib (AKTi) for 72 h. n.s.: non-significant, \**P* < 0.05; *t*-test.

Next, we focused on sgRNAs enriched in GNF-7-treated cells and thereby related to mutations conferring resistance to the drug (**Fig. 3E**). The presence of P53 in this list further highlighted the role of the G1/S transition. In addition, there was an enrichment for factors related to the suppression of PI3K/AKT signaling such as PTEN, TSC2 or TSC1 (**Fig. 3E, F**). In agreement with the screen data, CRISPR-mediated depletion of PTEN or TSC2 reduced the toxic effects of GNF-7 in HT-1080 cells (**Fig. S3G-I**). Furthermore, the senolytic effects of GNF-7 were potentiated through combination with the clinically approved AKT inhibitor capivasertib (**Fig. 3G**). In summary, this genetic screen confirmed the preferential toxicity of GNF-7 for growth arrested cells and identified strategies that can potentiate its senolytic activity.

### A GCN2-dependent ISR mediates the senolytic effects of GNF-7

To investigate the mechanism of action of the senolytic effects of GNF-7, we first conducted RNA sequencing (RNAseq) on senescent HT-1080^T^ cells exposed to GNF-7 for 6h in order to focus on the initial response to the drug. The comparison of proliferating and dox-treated HT-1080^T^ cells, was consistent with the onset of senescence, evidenced by a general downregulation of cell growth-related pathways together with the presence of several hallmarks from senescent cells such as lysosomal activity and P53-or TNF-dependent signaling (**Fig S4**). In what regards to the effects of the drug, the pathways upregulated by GNF-7 were primarily related to aminoacid availability and the integrated stress response (ISR) (**Fig. 4A**). Of note, from the 4 kinases of the ISR, the activation of the ISR by a deficit in aminoacids is triggered by the GCN2 kinase^31^. Consistently, western blot (WB) analyses revealed that GNF-7 triggered GCN2 phosphorylation, together with the activation of the ISR-dependent apoptotic factor CHOP, with this signaling being more acute in senescent cells than in proliferating ones (**Fig 4B**).

**Figure 4.**
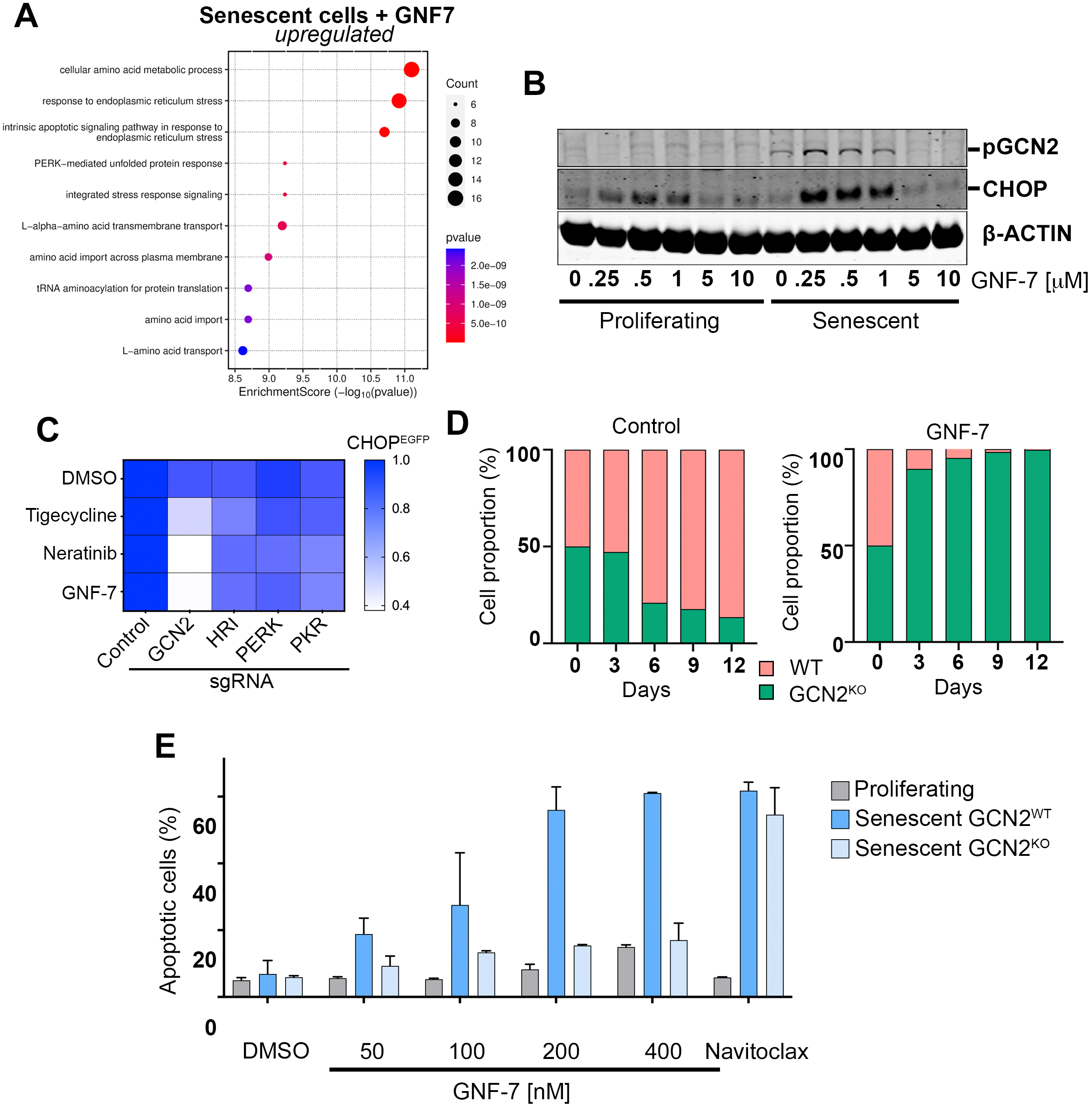
GCN2 activation mediates the senolytic effects of GNF-7. (**A**) Dot plot representing significantly upregulated biological processes in senescent HT-1080^T^ cells upon treatment with GNF-7 (200nM, 6 h). The X-axis indicates the enrichment score (-Log10pvalue). For the analysis the top 100 hits sorted by False Discovery Rate (FDR) and with a positive log2FoldChange were selected. Senescence was induced by exposure to doxycycline (1μg/ml) for 6 days. (**B**) WB analysis of phosphorylated GCN2 (pGCN2) and CHOP levels in proliferating and senescent HT’1080 cells. GNF-7 was used at the indicated concentrations for 3 h. Senescence was induced by palbociclib (5 μM) for 7 days. (**C**) Heatmap representing the EGFP median fluorescence intensity in DLD-1-CHOP^EGFP^ cells after treatment with the indicated drugs for 24 h. Cells were infected with mCherry-expressing lentiviral vectors carrying sgRNAs to target the four ISR kinases. Control cells were infected with the empty vector. EGFP MFI ratios between mCherry-positive (KO) and negative (WT) cells were calculated and normalized to the control levels for each treatment: untreated, Tigecycline 10 μM, Neratinib 500 nM and GNF-7 200 nM. (**D**) Competition assay in wild type (red, WT) and GCN2-deficient (green, GCN2^KO^) HT-1080 cells cultured for 12 days in the presence or absence of 50 nM GNF-7. **(E)** Percentage of apoptotic cells (measured by FACS as Annexin V positive and PI negative) upon GNF-7 treatment in proliferating cells, as well as senescent WT and GCN2^KO^ HT-1080 cells. Senescence was induced by palbociclib (5 μM) for 7 days. GNF-7 was used at the indicated doses, and navitoclax at 5 μM.

To further determine the ISR kinase responsible for triggering CHOP expression in response to GNF-7, we used a DLD-1 colorectal cancer line harboring a fluorescent reporter of the *CHOP* promoter (CHOP^EGFP^). As previously reported, GCN2 selectively mediated CHOP expression triggered by the antibiotic Tigecycline or the Tyrosine kinase inhibitor neratinib^27^ (**Fig. 4C** and **S5A**). Likewise, CRISPR mediated depletion of GCN2, but not PERK, PKR or HRI, abolished GNF-7-dependent CHOP expression (**Fig. 4C** and **S5B**). In agreement with our findings with CHOP in DLD-1 cells, GCN2 deletion abrogated the nuclear accumulation of ATF4 triggered by GNF-7 in HT-1080 cells (**Fig. S5C**).

Next, to determine the contribution of the activation of the ISR to the cellular effects of GNF-7, we first performed a cellular competition assay whereby parental HT-1080 cells expressing mCherry were mixed at a 1:1 ratio with GCN2-deficient HT-1080 cells expressing EGFP and cultured in the absence or presence of the drug. In control conditions, GCN2-deficient cells were progressively overgrown by parental cells, reflecting a lower fitness of the mutant population. However, this dynamic was strikingly reverted in the presence of GNF-7, reflecting a significant resistance to the compound in GCN2-deficient cells (**Fig. 4D**). Importantly, GCN2 deficiency rescued the apoptosis triggered by GNF-7 in senescent cells, demonstrating that GCN2 activation is responsible for the senolytic effects of the compound (**Fig. 4E**). Of note, GCN2-deficiency did not revert the senolytic effects of navitoclax, arguing against a general resistance of GCN2-deficient cells to senolytic compounds (**Fig. 4E**). Time-lapse videos measuring Caspase-3/7 activity confirmed the role of GCN2 in mediating the apoptosis triggered by GNF-7 in senescent cells (**Videos S3, S4**).

Recent works from us and others found that some multitarget TK inhibitors (TKi) activate GCN2 and target cancer cells with increased resistance to chemotherapy^27,32–34^. We thus explored whether the senolytic properties of GNF-7 were also observed with other TKi-s, that might share a common effect on GCN2. Supporting this view, we tested 6 different TKis and found that only those that activated the ISR (as measured by the nuclear translocation of ATF4) were senolytic for HT-1080 cells in a GCN2-dependent manner (**Fig. S5D, E**). Together, these experiments reveal that the senolytic effects of TKis such as GNF-7 are due to the activation of a GCN2-dependent apoptotic response.

### GNF-7 activates the kinase domain of GCN2

GCN2 is activated in cells by uncharged tRNAs that accumulate during amino-acid starvation, in a manner that depends on interaction with its binding partner GCN1 through the C-terminal histidyl-rRNA synthase domain of GCN2. Surprisingly, whereas, as previously reported, GCN1 deletion abolished the activation of the ISR induced by Tigecycline^27^, it did not affect the ISR-activation triggered by GNF-7 (**Fig. 5A**). Moreover, and in contrast to the effects of GCN2, GCN1 deletion did not rescue the senolytic effects of GNF7 in HT-1080 cells (**Fig. 5B**). These results indicate that the GNF-7-dependent activation of GCN2 occurs through a non-canonical mechanism and suggest that the compound could have a direct effect on GCN2.

**Figure 5.**
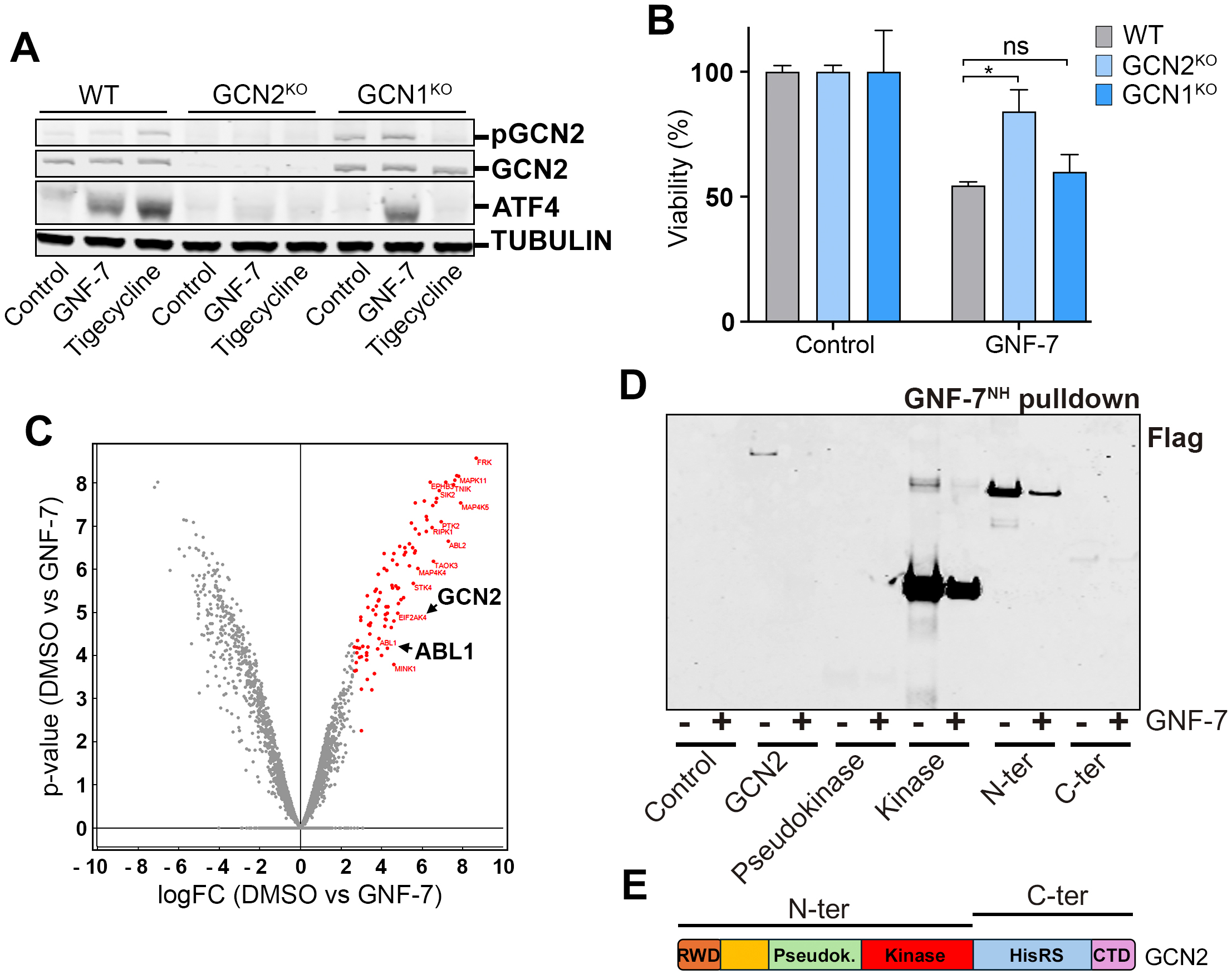
GNF-7 is an allosteric activator of GCN2. **(A)** WB analysis of GCN2, pGCN2, ATF4 and p-EIF2α levels in senescent WT, GCN2^KO^ or GCN1^KO^ HT-1080 cells after treatment with 200 nM GNF-7, 10 μM tigecycline or DMSO as a control for 16 h. TUBULIN levels are shown as a loading control. Senescence was induced by palbociclib (5 μM) for 7 days. **(B)** Normalized viability, as measured by CellTiter-Glo luminescent assay, from the experiment described in (A). **(C)** Proteomic analysis of GNF7^NH^ interactors (significant interactors are highlited in red) in HT-1080 protein lysates. The comparison is performed against samples prepared in the presence of an excess of parental GNF-7. **(D)** GNF-7^NH^ pull-down in GCN2-deficient HEK293T cells transfected with the indicated GCN2 expression plasmids. The WB shows the expression levels of the indicated constructs (based on Flag), showing both the control condition and the competition with an excess of the unmodified GNF-7. **(D)** Scheme of the GCN2 domain structure, illustrating the regions that were included in the N-ter and C-ter constructs. Sequence details for all cloned regions are provided in the methods section. n.s.: non-significant, \**P* < 0.05; *t*-test.

To determine whether GNF-7 binds to GCN2, we generated an amine-derivative of GNF-7 (GNF-7^NH^) and verified that the compound maintained its senolytic properties (**Fig. S6A, B**). Affinity pull-downs using NHS-Sepharose beads coupled to GNF-7^NH^ revealed that, in cell extracts, the compound bound its original target ABL1, as well as GCN2. These interactions were specific, as they were abrogated in the presence of an excess of parental GNF-7 (**Fig. S6C**). Using this approach, we next performed a proteome-wide characterization of GNF-7^NH^ interactors. This analysis identified 101 proteins that were significantly enriched as binders of GNF-7^NH^, 87 of which were kinases (86%), supporting the robustness of our purification pipeline (**Fig. 5C** and **Table S3**). Of note, the rest of the interactors were proteins known to bind kinases such as FGF2, EIF5 or GSKIP, an observation that had been previously made in a systematic proteomic analysis of factors bound by clinically used kinase inhibitors^35^. GNF-7 bound multiple kinases from the TK or MAPKK (STE) families but had a distinct effect in binding PERK and GCN2 and no other members of this family of kinases (**Fig. S6D**).

To define the protein domain in GCN2 mediating the interaction with GNF-7, we expressed different domains of GCN2 in a GCN2-deficient background and repeated the affinity pulldown with GNF-7^NH^. These experiments revealed that GNF-7 bound the kinase domain of GCN2, but not its pseudokinase or C-terminal histidyl-tRNA synthetase domains (**Fig**. **5D, E**), suggesting that the stimulatory effect of GNF-7 occurs through a direct effect of the drug on the GCN2 kinase domain.

To explore how the activation might occur at the molecular level, we performed docking experiments using RoseTTAFold All-Atom (RFAA). These analyses revealed that the TKIs that had an agonistic effect on GCN2 occupied its ATP-binding pocket as well as an adjacent allosteric pocket that is present in the kinase domain. In contrast, other TKis that fail to activate GCN2 bound exclusively to its ATP-binding pocket (**Fig. 6A**). Based on this information, we synthesized several GNF-7 derivatives (**Fig. 6B**). Eliminating the fragment of GNF-7 that occupies the allosteric pocket abrogated GCN2 activation (cpd. 1), an observation that was also seen with other changes to this part of the molecule (cpds. 4,5) (**Fig. 6C, S7**). Significantly, substitutions that retained the capacity to activate GCN2 were senolytic (cpds. 2, 3, 6, 7), whereas those that failed to activate GCN2 were not (**Fig. 6D**).

**Figure 6.**
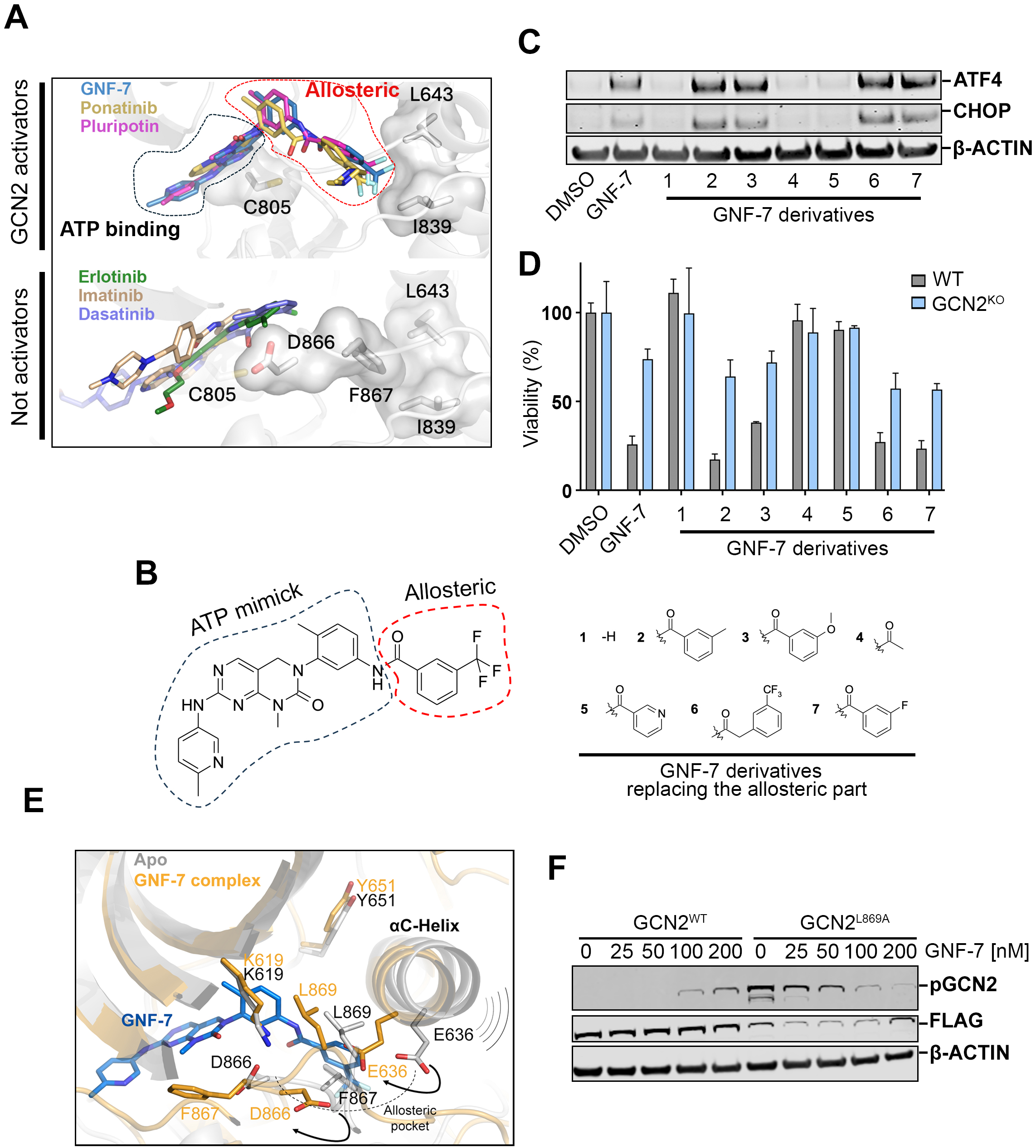
Senolytic TKIs bind to an allosteric pocket on the GCN2 kinase domain. (**A**) Predicted model for the binding of the GCN2 kinase domain to TKIs that activate GCN2 (GNF-7, Ponatinib and pluripotin) or fail to activate it (Erlotinib, Imatinib and Dasatinib). Highlighted areas illustrate the regions used for ATP-binding (black dashed line) or the adjacent allosteric pocket (red dashed line). Docking analyses were performed using RoseTTAFold All-Atom (RFAA). (**B**) Chemical structures of GNF-7 and the modified derivatives replacing the allosteric part. Highlighted areas illustrate the regions of the molecule predicted to bind to the ATP-binding or allosteric pockets. (**C**) WB analysis of ATF4 and CHOP levels in HT-1080 cells treated with 400nM of GNF-7 or the indicated derivatives for 6 h. β-ACTIN was employed as a loading control. (**D**) Normalized viability, as measured by CellTiter-Glo, of senescent WT and GCN2^KO^ HT-1080 cells treated with with 400nM of GNF-7 or the modified derivatives for 72 h. Senescence was induced by palbociclib (5 μM) for 7 days. (**E**) Predicted model of the kinase domain of GCN2 not bound to any ligand (Apo form, grey), or bound to GNF-7 (yellow), illustrating the position of L869 and its potential displacement by GNF-7. (**F**) WB analysis of pGCN2 and FLAG levels in GCN2-deficient HEK293T cells, transfected with constructs expressing wild type GCN2 (GCN2^WT^) or the L839 mutant (GCN2^L869A^), and treated with increasing concentrations of GNF-7. β-ACTIN was employed as a loading control.

As to how this interaction could promote GCN2 activation, we noted that the region of GNF-7 that occupies the allosteric pocket could potentially interact with L869 (**Fig. 6E**). This is relevant since previous work in yeast had shown that this Leu limits GCN2 catalytic activity by constraining the rotation of the adjacent helix αC, so that yeast L856A mutants were hyperactive^36^. On this basis, we hypothesized that the activation of GCN2 by GNF-7 could be mediated by an effect of the drug displacing L869, thereby mimicking the effect of the mutation. This mechanistic model implies that a L869A mutant should be (a) constitutively active, (b) not activatable by GNF-7 and (c) still inhibited by GNF-7 since the molecule would still interfere with ATP binding. Significantly, all of these predictions were confirmed when expressing the L869A mutant in GCN2-deficient cells (**Fig. 6F**). Together, these results support a model where the activation of GCN2 by TKis such as GNF-7 occurs through the direct binding of these compounds to an allosteric pocket that is present in the GCN2 kinase domain, which displaces inhibitory hydrophobic residues that limit the catalytic cycle of the kinase.

### A one-two-punch strategy with GNF-7 targets senescent cells in vivo

Our initial objective was to identify compounds that can target cancer cells that have resisted an initial chemotherapy. To test if this was the case with GNF-7, we first performed an allograft of B6-F10 mouse melanoma cells, which were first treated with doxorubicin for 3 days, and subsequently with GNF-7. In this model, GNF-7 synergized with doxorubicin in reducing tumor sizes and prolonged the survival of mice (**Fig. S8**). Nevertheless, since doxorubicin induces significant apoptosis, we decided to test another model where the senolytic effects of GNF-7 could be better evaluated.

To this end, we used xenografts of HT-1080 cells, which were treated with palbociclib for 1 week to induce senescence, followed by treatment with GNF-7 (**Fig. 7A**). Strikingly, GNF-7 led to the regression of HT-1080 xenografts treated with palbociclib (**Fig. 7B, C**). Furthermore, a treatment with GNF-7 did not significantly affect the growth of the tumors when these were not previously treated with palbociclib, underscoring that the compound is selectively targeting palbociclib-treated cells. Immunohistochemical (IHC) analyses of the xenografts confirmed the senolytic effects of GNF-7, by the clearance of cells with high levels of p21 or SA-ý-Gal activity (**Fig 7D**). A clear visualization of the senolytic properties of GNF-7 came from the analysis of cleaved-caspase 3 activity (C3A). Whereas most chemotherapies target rapidly growing cancer cells, IHC analyses revealed an inverse correlation between KI67 and C3A levels in the xenografts that received the sequential treatment with palbociclib and GNF-7 (**Fig. 7E**). In summary, these experiments provided solid proof of the senolytic effects of GNF-7 *in vivo*. In addition, and regardless of senescence, these data exemplify that GNF-7 greatly potentiates the anticancer effects of CDK4/6 inhibitors, when applied in a one-two-punch regimen.

**Figure 7.**
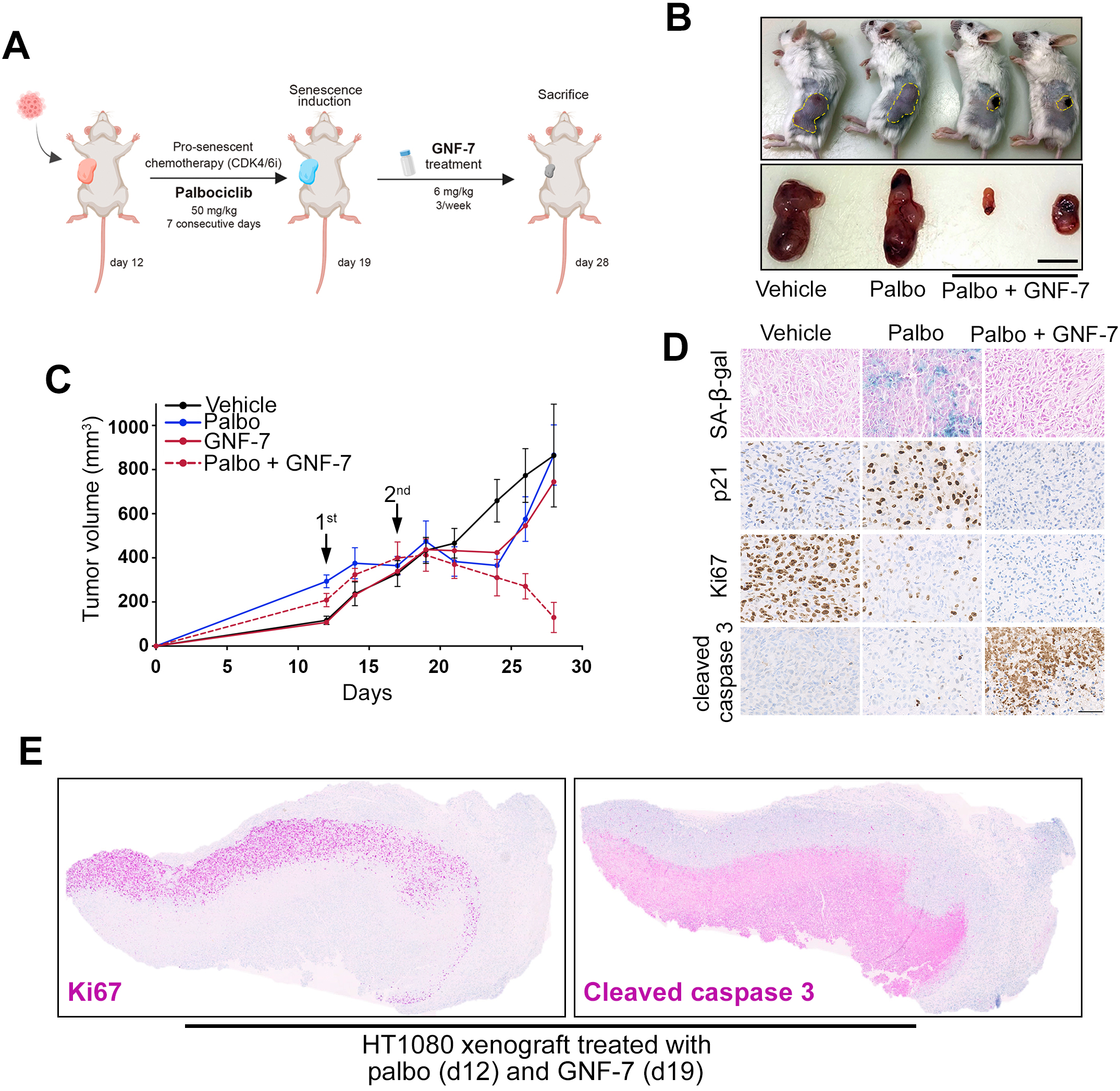
A one-two-punch therapy with GNF-7 targets palbociclib-treated senescent cells *in vivo*. (**A**) Schematic representation of the xenograft experiment. Human HT-1080 cells were injected into the flank of SCID mice. Palbociclib was administered for 7 consecutive days (50mg/kg, oral gavage), after which mice were treated with GNF-7 three times a week for 9 days (6mg/kg, oral gavage). Mice were sacrificed on day 28. (**B**) Representative images of mice and isolated tumors at the endpoint of the experiment (day 28). Scale bar (black) represents 1 cm. (**C**) Tumor growth curves (mm3) of HT-1080 xenografts treated with palbociclib or GNF-7 alone, or with GNF-7 being applied after the initial palbociclib treatment. Arrows indicate the starting point for palbociclib (1^st^, day 12) and GNF-7 treatments (2^nd^, day 19). (**D**) Immunohistochemical analysis of SA-β-gal, p21, KI-67 and cleaved caspase-3 expression in xenografts isolated from the experiment defined in (A). Scale bar (black) indicates 50 μm. (**E**) KI-67 and cleaved caspase-3 expression in whole HT-1080 tumor xenografts treated with palbociclib followed by GNF-7.

## Discussion

We here present our discovery of the multitarget TKi GNF-7 as a novel senolytic drug. An interesting aspect of this work is our observation that the drug is preferentially toxic for G1-arrested cells, before senescence is established. In fact, our primary screening was conducted in cells where the DDR was activated, but where senescence was not yet in place. Similar observations were made with dasatinib, suggesting that senolytic agents might be exploiting a vulnerability of rapidly growing cancer cells that are suddenly arrested in G1. As to what this vulnerability is, this remains to be better understood. One possibility is that rapidly proliferating cells that are growth arrested, maintain high anabolic rates due to the persistence of PI3/mTOR signaling, creating metabolic distress and issues with autophagic clearance of damaged organelles^37^. Of note, arresting cells in G1 is different from the G0 state, which refers to cells outside of the cell cycle. This explains why GNF-7 does not have widespread toxicity in mice, as most cells in an animal remain quiescent in G0.

On another note, our data further support that the mechanism of action of drugs, even those in oncological practice, often involves targets different to the originally intended ones^38^. This is particularly true for kinase inhibitors, which are often designed to occupy the ATP-binding pocket which has significant similarities among kinase domains. Systematic proteomic analysis of interactors of kinase inhibitors confirmed the widespread knowledge that these drugs bind to multiple kinases in cells, even when these medicines are routinely administered as specific inhibitors of a single target^35^. To add to this theme, the work presented here, together with previous manuscripts including our own work^27,32–34^, reveals that the toxicity of certain TKis might be surprisingly unrelated to TKs, and partly due to an effect of these compounds in mediating the activation of the Ser/Thr-kinase GCN2. Here, we add to this theme by identifying that this effect is preferentially toxic for senescent cells and by unraveling the mechanism that clarifies the structural determinants of this phenomenon. We hope that our discoveries can be used to guide the rational design of novel, ad-hoc, GCN2 agonists as a new class of anticancer agents.

To end, our research confirms that the use of senolytic drugs is maximized in the context of “one-two-punch” regimens^19^. While such an approach is often not prioritized in clinical trials, which rather focus on simultaneous drug combinations or in administering the second drug only after patients have relapsed from the initial treatment, it seems evident that the efficacy of these therapies is notably improved when sequentially applied after a pro-senescent chemotherapy. Whether the senolytic effects of GNF-7 can be beneficial in other age-related pathologies remains to be tested. To the very least, and regardless of its potential as a broad-spectrum senolytic agent, our work describes that GNF-7 can be used to maximize the clearance of cancer cells in tumors that have been previously treated with cytostatic treatments such as hormonotherapy in ER+ breast cancer or CDK4/6 inhibitors.

## Materials and Methods

### Cell Culture

All cell lines were incubated at 37 °C, in a humidified air atmosphere with 5% CO2. Human cell lines (HT-1080, MCF-7, DLD-1, RPE, HEK293T) and B16-F10 murine cancer cells were cultured using standard high glucose Dulbecco’s Modified Eagle Medium (DMEM, Sigma, D5796) supplemented with 10% Fetal Bovine Serum (FBS, Sigma) and 100U/ml Penicillin/Streptomycin (Pen/Strept, Gibco, 11548876). Mouse embryonic stem cells (mES) were grown on gelatin and feeder layers in high glucose DMEM supplemented with 15% knockout serum replacement (Invitrogen), Leukemia Inhibitory Factor (LIF, 1000U/ml), 0.1mM non-essential amino acids, 1% glutamax, and 55mM β-mercaptoethanol. Early passaged mouse embryonic fibroblasts (MEF) were irradiated with 80 Gray to be used as feeder layers. CCNE1 overexpressing RPE cells (RPE hTERTCas9 TP53-/-CCNE1OE) were kindly provided by Dr. Daniel Durocher^39^.

Patient-derived ER+ breast cancer organoids were kindly supplied by Dr. Miguel Quintela’s laboratory at CNIO. They were trypsinized using TrypLE Express (Gibco, 12604013), and the resulting pellets were washed in 10mL of Ad-DF+++ medium and centrifuged at 1200rpm for 5 minutes. Pellets were then resuspended in 45µl drops of cold Matrigel Matrix (Corning, CLS354234) and plated in prewarmed 24-well culture plates, allowing solidification at 37°C for 1 hr. Once solidified, 500µl of complete organoid medium was added to the wells and plates were incubated at 37 °C in a humidified air atmosphere with 5% CO2. Culture medium was refreshed every 4 days and organoids were passaged upon necessity. For 96-well culture plates, organoids were embedded in 1:1 mixture of Matrigel Matrix and medium. For cryopreservation, organoids were frozen in Stem Cell Freezing Media (ATCC, ACS-3020) and stored at -80°C. The composition of the media used for organoid cultures, as well as a full list of all the reagents used in this study, is provided at **Table S4**).

### CRISPR editing

To generate gene knockouts, single-guide RNAs (sgRNAs) were selected using the CRISPOR tool (Concordet JP. and Haeussler M., Nucleic Acids Research, 2018; https://crispor.gi.ucsc.edu/crispor.py) or the sgRNAs oligonucleotide sequences from the Human CRISPR Knockout Pooled Library Gattinara (Addgene, Pooled Library #136986)^40^. Repair ability was predicted using inDelphi (https://indelphi.giffordlab.mit.edu/)^41^. sgRNA sequences were cloned into the following lentiviral vectors: plentiCRISPR v2-Blast (Plasmid #83480), plentiCRISPRv2-puro (Plasmid #52961), pLentiGuide-mCherry and pLentiGuide-GFP (kind gift of Cristina Mayor-Ruiz) using the NEB® Golden Gate Assembly Kit (BsmBI-v2) (NEB, 174E1602S), following manufacturer’s instructions. All sgRNA sequences are available in **Table S4**.

### CRISPR/Cas9 screen

A clonal mESC-Cas9 cell line was transduced with the Mouse Toronto Knockout (mTKO) CRISPR Library (Addgene 159393, two-vector system)^26^, which contains 94,528 single-guide RNAs targeting 19,463 murine genes (∼5 gRNAs/gene). A total of 65 million mESC cells were transduced at a low MOI (0.3), maintaining 200-fold library coverage. Infected cells were treated with with GNF-7 (200 and 400nM) and with DMSO as a control. Uninfected control cells were also seeded in parallel. Resistant cell pools were then harvested and stored at -80°C on day 10. DNA from resistant pools was extracted using Gentra Puregene Blood Kit (Qiagen, 158445), following manufacturer’s instructions. Then, to identify sgRNAs, the U6-sgRNA cassette was amplified by PCR using the KAPA HIFI Hot Start PCR kit (Roche, KK2502). The PCR product was precipitated with 3M sodium acetate in 100% ethanol at -80 °C for at least 20 minutes. Samples were purified either in agarose gel or using magnetic beads and sent to Illumina sequencing for sgRNA detection.

Data analysis was performed using the Model-based Analysis of Genome-wide CRISPR/Cas9 Knockout (MAGeCK) pipeline to identify relative sgRNA abundances^30^. Raw reads were normalized to total sgRNAs counts to account for library size effect. Sequencing reads from fastq files were then mapped to the sgRNA reference library, and sgRNA counts were calculated using a negative binomial model. To identify both positively selected genes –whose sgRNAs are enriched– and negatively selected ones –with depleted sgRNAs–, sgRNAs were ranked based on their enrichment p-values, using the alpha-Robust Rank Aggregation (α-RRA) statistical method.

### Lentiviral production

HEK293T cells were co-transfected with a lentiviral transfer vector and third-generation packaging plasmids (pMDL, pRev and pVSV-G), using Lipofectamine 2000 (ThermoFisher #11668019) and Opti-MEM Reduced Serum Medium (Gibco, 31985070), following manufacturer’s instructions. The viral supernatant was collected 48-72 h post-transfection, filtered through a 0.45µm filter, added to cells with 6mg/mL polybrene and incubated for 8 hr. After 48 hr, infected cells were selected using puromycin (2µg/mL, Sigma, P8833) or blasticidin (10µg/mL, Life Technologies, A11139-03).

### Flow cytometry

To analyze cell cycle profiles, cell pellets were fixed dropwise with cold ethanol (70% v/v) while vortexing, followed by an incubation on ice for 30 minutes. Samples were then either stored at -20°C or immediately washed twice in PBS and centrifuged at 2000rpm for further processing. Cells were treated with 100μg/mL RNase (Qiagen) and 100μg/mL propidium iodide (PI, Sigma-Aldrich). Samples were analyzed using the FACS Canto II (BD Biosciences). Flow cytometry data was analyzed using FlowJo software.

### Competition assays

eGFP-expressing cells harboring a gene-specific sgRNA were mixed in a 1:1 ratio (750,000 cells of each condition) with mCherry-expressing control cells. On the following day, cells were treated either with 50nM GNF-7 or DMSO as a control. Cells were passaged and analyzed twice a week by flow cytometry. Cells were trypsinized and resuspended in PBS with 4’,6 diamidino 2 phenylindole (DAPI) to stain viable cells, and the different cell populations were subsequently analyzed using the BD FortessaTM flow cytometer (BD Biosciences).

### Cell viability

For High-throughput microscopy (HTM)-based quantification, cells were seeded in μCLEAR bottom 96-well plates (Greiner Bio-One) and treated with the indicated concentrations of drugs. After 48-72 hr, cells were fixed with 4% Paraformaldehyde (PFA) for 10 minutes, permeabilized with 0.5% Triton X-100 for 10 minutes and stained with DAPI. Then, HTM images were acquired using an Opera Phenix™ Plus High Content Imaging System (Perkin Elmer) or an ImageXpress Pico Automated Cell Imaging System (Molecular Devices). Generally, images were taken using the 20X magnification lens, and nuclei were counted to measure cell viability. For luminescence-based quantification, cell viability was assessed using CellTiter Glo Assay (CTG, Promega), following manufactureŕs instructions.

### Immunofluorescence and high throughput microscopy

For immunofluorescence, cells were fixed with 4% PFA prepared in PBS buffer and permeabilized with 0.5% Triton X-100 after fixation. Blocking was performed for 30 minutes using a solution of 3% Bovine serum albumin (BSA) and 0.1% Tween-20 in PBS. To monitor CHOP levels, we used the lentiviral plasmid pCLX-CHOP-dGFP (Addgene, 71299). For high throughput microscopy, cells were grown on µCLEAR bottom 96-well plates (Greiner Bio-One) and immunofluorescence of ATF4 (Cell Signaling, 11815) was performed using standard procedures. Analysis of DNA replication by EdU and transcription by were done using Click-It kits (Invitrogen) following the manufacturer’s instructions. In all cases, images were automatically acquired from each well using an Opera High-Content Screening System (Perkin Elmer). A 20x magnification lens was used and images were taken at non-saturating conditions. Images were segmented using DAPI signals to generate masks matching cell nuclei, from which the mean signals for the rest of the stainings were calculated. Data representation was performed using GraphPad Prism 9 software.

### Western Blotting

Cells were harvested and lysed at 4°C on a shaker for 20 minutes using Urea buffer (50 mM Tris, pH 7.5, 8 M urea, and 1% CHAPS) supplemented with protease and phosphatase inhibitors (Sigma Aldrich). For other samples, RIPA buffer (Tris HCl 50 mM, pH 7 .4, NP 40 1%, Na deoxycholate 0.25%, NaCl: 150 mM, EDTA 1 mM) was employed as an alternative lysis buffer, with an incubation step of 30 minutes at 4°C. After lysis, the protein fraction was isolated by centrifugation at 13,200rpm for 10 minutes at 4°C and protein concentrations were determined by the Bradford Protein Assay (Bio-Rad). Protein extracts were mixed with NuPAGE LDS (LifeTechnologies) supplemented with 10 mM dithiothreitol (DTT, Sigma) and were incubated at 99 °C for 10 minutes. Samples were resolved in precast 4-16% gradient SDS-polyacrylamide gels (Invitrogen) and transferred following standard Western blot protocols. After blocking, the membrane was incubated overnight at 4 °C with primary antibodies (**Table S4**). The following day, membranes were incubated with a fluorophore-conjugated secondary antibody at RT, and protein detection was performed using the Li-Cor LCx system (Biosciences) to detect the protein.

### Annexin staining

Cells were treated with GNF-7 and pan Caspase Inhibitor Z-VAD-FMK (Bio-techne R&D SYSTEMS S.L, FMK001) for 48 hr. Cells were collected and stained with APC Annexin V (550474, BD Biosciences), following manufacturer’s instructions and using propidium iodide (PI) staining to select viable cells. Apoptotic cells were identified by flow cytometry, using the FACS Canto II (BD Biosciences), and data was analyzed using FlowJo software.

### Senescence

To induce senescence, different drugs were employed according to the cell line, as described in the manuscript. In HT-1080^T^ cells, telomere uncapping-induced senescence was triggered by 1μg/mL doxycycline treatment during 6 consecutive days. Senescence was also induced using 5μM palbociclib for 7 days or 50-100nM doxorubicin treatment for 5 days. To detect SA-β-gal activity, cells were fixed and stained with the senescence-associated β-galactosidase kit (Cell Signaling, 9860), following the manufacturer’s protocol.

### Cell cycle arrest

To arrest cells in G1, these were subjected to a double thymidine block. To do so, cells were first treated for 16 h with 2.5mM thymidine (Sigma, T-1895), released for 8 h in thymidine-free media, and blocked again with 2.5mM of thymidine for 16 hr.

### RNA-seq

Total RNA was extracted from cells using the Absolutely RNA Microprep kit (Agilent, 400805), following manufacturer’s instructions.. The sequencing library was constructed with the QuantSeq 3’ mRNA-Seq Library Prep Kit (Lexogen), and approximately 10 million reads were obtained by Illumina sequencing. Differential expression analysis was conducted using Salmon software (v1.3.2) for pseudoalignment of the fastq files, using the Ensembl human genome version (Homo_sapiens. GRCh38). Output files were analyzed in R Studio with DESeq2, and deregulated genes were visualized using R Studio’s plotting functions.

### Chemical affinity purification

Affinity purification pull-downs with GNF7^NH^ protocol was conducted as previously described^42^. Briefly, GNF7^NH^ as coupled to NHS-Activated Sepharose 4 Fast Flow Beads (GE LifeSciences, 17090601). For each condition, 100μL of NHS-sepharose beads were washed three times with 500μL of Dimethyl sulfoxide (DMSO), centrifuging at 800rpm for 3 minutes at RT. Beads were then resuspended in 50μL of DMSO, and 2.5µl of 10mM GNF-7-NH along with 0.75μL of triethylamine (TEA, Sigma, T0886) were added to each sample. Following an incubation of 16-24 h on a roto-shaker at RT, 2.5μL of ethanolamine (Sigma-Aldrich, 11016-7) were added to block any remaining free amino groups on the beads, followed by at least 8 h of incubation on a roto-shaker at RT. After blocking, beads were centrifuged and washed twice with 500μL of DMSO before proceeding with the pull-down assay.

To prepare cell lysates, HT-1080 cells were resuspended in lysis buffer (50mM Tris pH 7.5, 0.2% NP-40, 5% glycerol, 1.5μM MgCl2, 100μM NaCl, 1μM EDTA), using 500μL of lysis buffer per 100 million cells. Buffer was further supplemented with protease inhibitors and benzonase (Merck, 70746) and incubated on ice for 30 minutes. To improve the lysis, cells were passed through a syringe multiple times. The protein fraction was then isolated by centrifugation at 12,000 g for 30 min at 4°C, and protein concentration was determined using the BCA assay (Fisher Scientific, 23225). A total of 4.5 mg of protein lysate was pre-treated with either DMSO or 100μM of parental GNF-7 on a roto-shaker for 1 hr.

To conduct protein affinity purification and elution, drug-coupled beads were first washed three times with lysis buffer by centrifugation. The pretreated cell lysates were then gently resuspended into the beads and incubated for 2 h on a roto-shaker at 4°C. After incubation, the beads were washed three times with lysis buffer at 4 °C. The beads were then resuspended in washing buffer (50μM pH8 HEPES, 150μM NaCl and 5μM EDTA) and transferred to unplugged mini Bio-Spin Chromatography Columns (BioRad, 7326207), which had been previously equilibrated in the same buffer, and washed three times with washing buffer by centrifugation (200g for 1 minute at 4°C). Subsequently, columns were plugged and transferred to RT, and beads were incubated with elution buffer (washing buffer supplemented with 4% SDS) for 15 minutes. To eluate the proteins, columns were unplugged, placed in 1.5 mL Eppendorf tubes and centrifuged for 1 minute at RT. Protein eluates were either stored at -80°C for proteomics analysis or further processed for Western blot, adding NuPAGE LDS plus DTT and boiling samples at 99 °C for 10 minutes.

### Proteomics

Proteins were reduced and alkylated (15 mM TCEP, 25 mM CAA, 2% SDS, 100 mM TEAB pH 7) 1h at 45 °C in the dark. Digestion was performed using the on-bead Protein Aggregation Capture (PAC) method^43^ with MagReSyn® Hydroxyl microparticles (ReSyn Biosciences) at a protein-to-bead ratio of 1:2, in an automated KingFisher instrument (Thermo Fisher Scientific). Proteins were digested for 16 h at 37 °C using a mix of trypsin (Sigma, sequencing grade, 0.2µg per sample) and LysC (Wako, 0.2µg per sample) in 50 mM TEAB (pH 8.0). After digestion, samples were acidified to a final concentration of 0.5% formic acid (FA).

LC-MS/MS was performed by coupling an UltiMate 3000 RSLCnano LC system to a Orbital Exploris 480 mass spectrometer (Thermo Fisher Scientific). Peptides were loaded into a trap column (Acclaim™ PepMap™ 100 C18 LC Columns 5 µm, 20 mm length) for 3 min at a flow rate of 10 µl/min in 0.1% FA. Then, peptides were transferred to an EASY-Spray PepMap RSLC C18 column (Thermo) (2 µm, 75 µm x 50 cm) operated at 45 °C and separated using a 60 min effective gradient (buffer A: 0.1% FA; buffer B: 100% ACN, 0.1% FA) at a constant flow rate of 250 nL/min. The gradient ranged from 2% to 6% of buffer B in 2 min, from 6% to 33% B in 58 minutes, from 33% to 45% in 2 minutes, ending with 10 additional minutes at 98% B. Peptides were sprayed at 1.5 kV into the mass spectrometer via the EASY-Spray source and the capillary temperature was set to 300 °C. The mass spectrometer was operated in a data-independent acquisition (DIA) mode using 60,000 precursor resolution and 15,000 fragment resolution. Ion peptides were fragmented using higher-energy collisional dissociation (HCD) with a normalized collision energy of 29. The normalized AGC target percentages were 300% for Full MS (maximum IT of 25 ms) and 1000% for DIA MS/MS (maximum IT of 22 msec). 8 m/z precursor isolation windows were used in a staggered-window pattern from 396.43 to 1004.70 m/z. A precursor spectrum was interspersed every 75 DIA spectra. The scan range of the precursor spectra was 390-1000mz.

Raw files were processed with DIA-NN (1.8.2) using the library-free setting against a human protein database (UniProtKB/Swiss-Prot, one protein sequence per gene, 20,614 sequences, downloaded in June, 2021) supplemented with contaminants. Precursor m/z range was set from 396 to 1010, and all other settings were set at their default values.

The protein group intensity table “report.pg_matrix.tsv” was loaded in Prostar 1.26.4^44^ for further statistical analysis. Briefly, proteins with two or less valid values in at least one experimental condition were filtered out. Missing values were imputed using the algorithms SLSA^45^ for partially observed values and DetQuantile for values missing on an entire condition. Differential analysis was done using the empirical Bayes statistics Limma. Proteins with a p.value < 0.01 and a log2 ratio DMSO vs Parent >2.65 were defined as potential interactors. The ratio threshold was determined by making an Empirical Cumulative Distribution Function using fold change absolute values and finding the first point where the slope is ≤ 0.05. The FDR was estimated to be below 1% by Benjamini-Hochberg. All pulldown conditions were analyzed in biological triplicates, and raw data for all replicates have been deposited in PRIDE (PXD065521).

### Structural analyses

Predicted models of the kinase domain structure and in complex with various ligands were generated by open-source deep learning tools. On one hand, the protein structure of the kinase domain of GCN2 (residues A573 to Q1005) and complexed with ATP were predicted using AlphaFold v3^46^. The structural models to predict ligand binding (kinase domain) were conducted using RoseTTAFold All-Atom (RFAA)^47^ and Boltz-1, an accurate open-source based on AF3 ^48^. Ligand binding predictions for GNF-7, ponatinib, pluripotin, erlotinib, imatinib and dasatinib were performed using RoseTTAFold All-Atom. Alternatively, predictions for GNF-7 derived compounds (ETP00058723, ETP00058730, ETP00058731, ETP00058735, ETP00058738 and ETP00058739) were executed using Boltz. The structural comparison between compounds and figures were performed using PyMOL 3.1^49^.

### Animal studies

For the allograft experiment, 5×10^5^ exponentially growing B16F10 cells were subcutaneously inoculated into the flank of immunocompetent syngeneic C57BL/6 mice acquired from Charles Rivers. Tumors were allowed to grow until reaching an initial volume of around 150-200 mm3 and mice were randomized into three groups: doxorubicin alone (4mg/kg three times a week, intraperitoneal administration), doxorubicin followed by GNF-7 (15mg/kg three times a week, oral gavage administration) and doxorubicin followed by navitoclax (50mg/kg every day, oral gavage). For the xenograft experiment, 10^6^ exponentially growing human fibrosarcoma HT-1080 cells were inoculated into the flank of 8-week severe combined immunodeficient (Prkdcscid) mice. In this case, mice were randomized into four groups: untreated, Palbociclib (50mg/kg for 7 consecutive days, oral gavage administration), GNF-7 (6mg/kg three times a week, oral gavage) and the combination of Palbociclib followed by GNF-7 treatment. In both scenarios, tumours were measured three times a week, using the standard formula (length x width x 0.5), and mice were sacrificed according to humane endpoint (1,600 mm3), following standard ethical procedures. Tumours were extracted for further processing and analyses. Mice were housed in the pathogen-free facilities of the Spanish National Cancer Research Centre (CNIO) under standard conditions. All animal experiments were performed according to the Guidelines for Humane Endpoints for Animals Used in Biomedical Research and under the supervision of the Ethics Committee for Animal Research of the “Instituto de Salud Carlos III”.

### Immunohistochemical analyses

Tissues were fixed in formalin overnight and then embedded in paraffin for further processing and sectioning at the Histopathology Unit at the CNIO. Sections of 2.5 µm were treated with citrate for antigen retrieval, and processed for immunohistochemistry with hematoxylin and eosin staining, or p21, KI-67 and c3-cleaved antibodies, following standard protocols. Sections were also stained with the senescence-associated β-galactosidase kit (Cell Signaling, 9860) following the manufacturer’s guidelines. Images were scanned and digitalized with a MIRAX system (Zeiss), and they were captured with a Leica DM2000 LED optical microscope (20x or 40x lens).

## Chemistry

### Synthesis of GNF-7^NH^

N-{4-Methyl-3-[1-methyl-2-oxo-7-(6-piperazin-1-yl-pyridin-3-ylamino)-1,4-dihydropyrimido[4,5-d]pyrimidin-3(2H)-yl]-phenyl}-3-trifluoromethyl benzamide. Compound GNF-7^NH^ was synthesised from N-(3-(7-Chloro-1-methyl-2-oxo-1,4-dihydropyrimido[4,5-d]-pyrimidin-3(2H)-yl)-4-methyl phenyl)-3-(trifluoromethyl) benzamide (cas 839708-52-0,140 mg, 1 eq) by reaction with 1-Boc-4-(5-aminopyridin-2-yl)piperazine (100 mg, 1.2 eq), Pd_2_dba_3_ (56 mg, 0.2 eq), XPhos (28 mg, 0.2 eq) and K_2_CO_3_ (125 mg, 3 eq) in 2 mL of i-propanol in a degassed sealed tube with Argon, and heated at 100 °C for 2 h. On cooling, the mixture was concentrated, and the residue was purified by flash column chromatography on silica gel (0-70% EtOAc/cHexane) to give the BOC protected derivative which was dissolved in MeOH (3 mL) and Amberlyst(R)-15 (∼200 mg) was added. The reaction mixture was shaked for 2.5 h. The resin was filtered off and then treated with 4 mL of a 7N solution of NH_3_-MeOH for 10 min and filtered. The filtrate was concentrated, and the resulting residue was purified by HPLC in reversed phase and then SCX-2 cartridge to give the expected product GNF-7^NH^ as a white solid (25 mg). LCMS (ESI) rt = 3.438 min, *m/z* 618.4 [M + H]^+^. ^1^H NMR (300 MHz, DMSO-*d_6_*) δ d 10.54 (s, 1H), 9.32 (s, 1H), 8.45 (s, 1H), 8.30 (s, 1H), 8.27 (d, J = 7.9 Hz, 1H), 8.09 (s, 1H), 7.98 (d, J = 7.9 Hz, 1H), 7.87 (d, J = 8.5 Hz, 1H), 7.81 (m, 2H), 7.64 (d, J = 8.3 Hz, 1H), 7.32 (d, J = 8.3 Hz, 1H), 6.79 (d, J = 9.2 Hz, 1H), 4.68 (d, J = 13.9 Hz, 1H), 4.50 (d, J = 13.9 Hz, 1H), 3.31 (m, 7H), 2.75 (m, 4H), 2.13 (s, 3H).

### Synthesis of compounds 2-7

The compounds were synthesized following similar previously reported synthetic procedures^50^. 3-Methyl-N-{4-methyl-3-[1-methyl-7-(6-methyl-pyridin-3-ylamino)-2-oxo-1,4-dihydro-2H-pyrimido[4,5-d]pyrimidin-3-yl]-phenyl}-benzamide (**2**). White solid. LCMS (ESI) rt = 3.438 min *m/z* 494.2 [M + H]^+^. ^1^H NMR (300 MHz, DMSO-*d6*) δ 10.27 (s, 1H), 9.66 (s, 1H), 8.79 (d, J = 2.5 Hz, 1H), 8.16 (s, 1H), 8.05 (dd, J = 8.4, 2.6 Hz, 1H), 7.82 (d, J = 2.1 Hz, 1H), 7.76 (sbroad, 1H), 7.73 (m, 1H), 7.62 (dd, J = 8.3, 2.1 Hz, 1H), 7.41 (dd, J = 4.9, 1.5 Hz, 2H), 7.29 (d, J = 8.5 Hz, 1H), 7.18 (d, J = 8.5 Hz, 1H), 4.71 (d, J = 14.4 Hz, 1H), 4.52 (d, J = 14.3 Hz, 1H), 3.34 (s, 3H), 2.40 (s, 6H), 2.13 (s, 3H). 3-Methoxy-N-{4-methyl-3-[1-methyl-7-(6-methyl-pyridin-3-ylamino)-2-oxo-1,4-dihydro-2H-pyrimido[4,5-d]pyrimidin-3-yl]-phenyl}-benzamide (**3**). Beige solid. LCMS (ESI) rt = 3.316min, *m/z* 510.0 [M + H]^+^. ^1^H NMR (300 MHz, DMSO-*d_6_*) δ 10.29 (s, 1H), 9.66 (s, 1H), 8.79 (d, J = 2.5 Hz, 1H), 8.16 (s, 1H), 8.05 (dd, J = 8.4, 2.6 Hz, 1H), 7.81 (d, J = 2.1 Hz, 1H), 7.63 (dd, J = 8.3, 2.1 Hz, 1H), 7.55-7.42 (m, 3H), 7.29 (d, J = 8.5 Hz, 1H), 7.21 – 7.11 (m, 2H), 4.71 (d, J = 14.4 Hz, 1H), 4.52 (d, J = 14.3 Hz, 1H), 3.84 (s, 3H), 3.34 (s, 3H), 2.40 (s, 3H), 2.13 (s, 3H).

N-{4-Methyl-3-[1-methyl-7-(6-methyl-pyridin-3-ylamino)-2-oxo-1,4-dihydro-2H-pyrimido[4,5-d]pyrimidin-3-yl]-phenyl}-acetamide (**4**). White solid. LCMS (ESI) rt = 2.53 min, *m/z* 418.2 [M + H]^+^. ^1^H NMR (300 MHz, DMSO-*d_6_*) δ 10.00 (s, 1H), 9.65 (s, 1H), 8.79 (d, J = 2.4 Hz, 1H), 8.14 (s, 1H), 8.05 (dd, J = 8.5, 2.6 Hz, 1H), 7.62 (d, J = 2.0 Hz, 1H), 7.37 (dd, J = 8.3, 2.0 Hz, 1H), 7.20 (m, 2H), 4.66 (d, J = 14.3 Hz, 1H), 4.47 (d, J = 14.2 Hz, 1H), 3.34 (s, 3H), 2.40 (s, 3H), 2.08 (s, 3H), 2.03 (s, 3H).

N-{4-Methyl-3-[1-methyl-7-(6-methyl-pyridin-3-ylamino)-2-oxo-1,4-dihydro-2H-pyrimido[4,5-d]pyrimidin-3-yl]-phenyl}-nicotinamide (**5**). Beige solid. LCMS (ESI) rt = 0.5 min, *m/z* 481.2 [M + H]^+^. ^1^H NMR (300 MHz, DMSO-*d_6_*) δ 10.52 (s, 1H), 9.66 (s, 1H), 9.11 (d, J = 1.8 Hz, 1H), 8.79 (d, J = 2.5 Hz, 1H), 8.77 (dd, J = 4.8, 1.5 Hz, 1H), 8.32 – 8.25 (m, 1H), 8.16 (s, 1H), 8.05 (dd, J = 8.5, 2.6 Hz, 1H), 7.81 (d, J = 2.0 Hz, 1H), 7.65 – 7.54 (m, 2H), 7.31 (d, J = 8.5 Hz, 1H), 7.18 (d, J = 8.5 Hz, 1H), 4.71 (d, J = 14.2 Hz, 1H), 4.52 (d, J = 14.3 Hz, 1H), 3.34 (s, 3H), 2.40 (s, 3H), 2.13 (s, 3H).

N-{4-Methyl-3-[1-methyl-7-(6-methyl-pyridin-3-ylamino)-2-oxo-1,4-dihydro-2H-pyrimido[4,5-d]pyrimidin-3-yl]-phenyl}-2-(3-trifluoromethyl-phenyl)-acetamide (**6**). White solid. LCMS (ESI) rt = 3.75 min, *m/z* 562.3 [M + H]^+^. ^1^H NMR (300 MHz, DMSO-*d_6_*) δ 10.31 (s, 1H), 9.65 (s, 1H), 8.78 (d, J = 2.5 Hz, 1H), 8.13 (s, 1H), 8.04 (dd, J = 8.5, 2.6 Hz, 1H), 7.70 (s, 1H), 7.67 – 7.53 (m, 4H), 7.39 (dd, J = 8.3, 2.0 Hz, 1H), 7.24 (d, J = 8.5 Hz, 1H), 7.17 (d, J = 8.5 Hz, 1H), 4.65 (d, J = 14.4 Hz, 1H), 4.47 (d, J = 14.2 Hz, 1H), 3.78 (s, 2H), 3.31 (s, 3H), 2.40 (s, 3H), 2.09 (s, 3H). 3-Fluoro-N-{4-methyl-3-[1-methyl-7-(6-methyl-pyridin-3-ylamino)-2-oxo-1,4-dihydro-2H-pyrimido[4,5-d]pyrimidin-3-yl]-phenyl}-benzamide (**7**). White solid. LCMS (ESI) rt = 3.29 min, *m/z* 498.2 [M + H]^+^. ^1^H NMR (300 MHz, DMSO-*d_6_*) δ 10.39 (s, 1H), 9.66 (s, 1H), 8.79 (d, J = 2.5 Hz, 1H), 8.16 (s, 1H), 8.05 (dd, J = 8.4, 2.7 Hz, 1H), 7.83-7.75 (m, 3H), 7.64 – 7.56 (m, 2H), 7.46 (m, 1H), 7.30 (d, J = 8.5 Hz, 1H), 7.18 (d, J = 8.5 Hz, 1H), 4.71 (d, J = 14.3 Hz, 1H), 4.52 (d, J = 14.3 Hz, 1H), 3.34 (s, 3H), 2.40 (s, 3H), 2.13 (s, 3H).

### Statistics

All statistical analyses were performed using Prism software (GraphPad Software) and statistical significance was determined where the p-value was <0.05 (*), <0.01 (**), <0.001 (***) and <0.0001 (****). Survival data was evaluated using Kaplan-Meier analyses.

## Data Availability

Mass spectrometry data have been deposited to the ProteomeXchange Consortium via the PRIDE partner repository [1] with the dataset identifier PXD065521. RNA-seq data are available at the GEO repository with accession number GSE301995.

## Supporting information

Supplementary Figures S1-7, and legends for supplementary videos and Tables

## Acknowledgments

The authors want to thank the Chemical Biology Consortium Sweden for their help with the chemical libraries; Dr. Daniel Durocher for providing the CCNE1 overexpressing RPE cells; Dr. Miguel Quintela for providing patient-derived organoids; Dr. Cristina Mayor for advice on the chemical affinity purification; Drs. Carmen Varela and Terry Tomakinian for their work on the synthesis of GNF-7 derivatives; as well as the proteomics, genomics, molecular imaging, confocal microscopy and transgenics units of the CNIO for their technical support in this study. Work done in the O.F-C. lab was supported by grants from the Spanish Ministry of Science, Innovation and Universities (PID2021-128722OB-I00, co-financed with European FEDER funds), the Spanish Association Against Cancer (AECC; PROYE20101FERN), Cancerfonden foundation (CAN 21/1529) and the Swedish Research Council (VR) (538-2014-31). The authors declare no competing financial interests.

## Author Contributions

G.L. contributed to most of the experiments; M.M. contributed to most of the experiments, their experimental design as well as the supervision of the project; W. A. conducted the initial screens and validation experiments; M.H. helped with screening experiments; M.L., A.G., J.M., B.N., S.F. and C.C. contributed to experiments related to the genetic screen and subsequent validations; E.J. and R.F. conducted the structural modelling experiments; E.Z. and M.I. conducted the proteomic analyses; S.M. and J.P. provided the chemical derivatives of GNF-7; M.E.A. and L.L. provided technical help; D.H. contributed to the experimental design and supervision of the project; O.F. conceived the experimental plan, coordinated the study, supervised the experiments and wrote the manuscript.

