## Supplementary Figures S1-7, and legends for supplementary videos and Tables for "The tyrosine kinase inhibitor GNF-7 targets senescent cells through allosteric activation of GCN2"

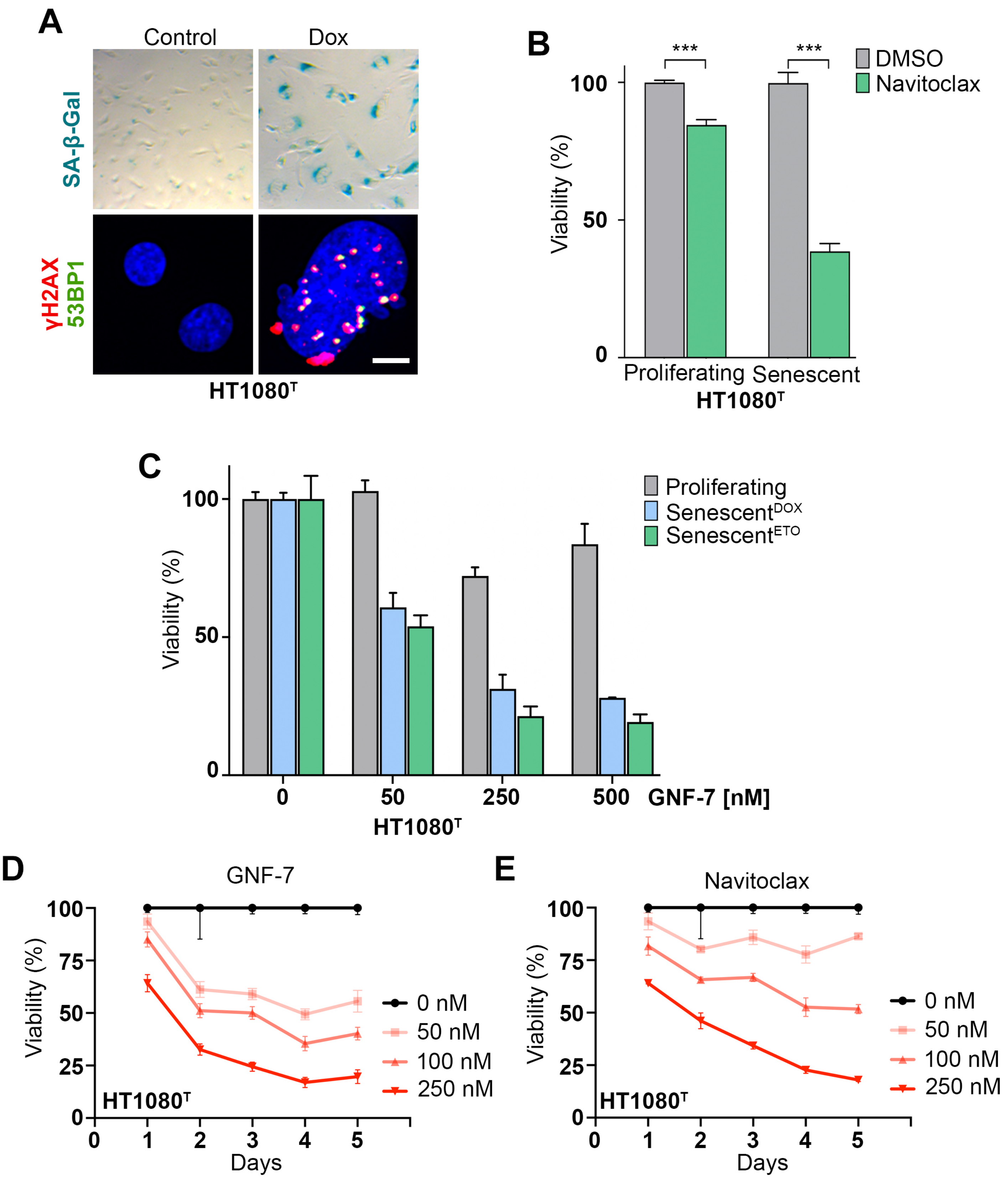
A
Control
Dox
SA-β-Gal
γH2AX
53BP1
HT1080T
B
***
***
DMSO
Navitoclax
Viability (%)
100
50
0
Proliferating
Senescent
HT1080T
C
Proliferating
SenescentDOX
SenescentETO
Viability (%)
100
50
0
0
50
250
500
GNF-7 [nM]
HT1080T
D
GNF-7
Viability (%)
100
75
50
25
0
0 nM
50 nM
100 nM
250 nM
HT1080T
0
1
2
3
4
5
Days
E
Navitoclax
Viability (%)
100
75
50
25
0
0 nM
50 nM
100 nM
250 nM
HT1080T
0
1
2
3
4
5
Days

**Figure S1. Senolytic effects of GNF-7 in different senescence models.** (**A**) Top: SA-β-gal assay in HT-1080$^{T}$ cells after 1 μg/mL doxycycline treatment for 6 days. Bottom: Representative immunofluorescence of γH2AX (red) and 53BP1 (green) foci in cells treated as for the top pannels. DAPI (blue) was employed to stain DNA. Scale bar (white) indicates 2μm. (**B**) Normalized viability, as measured by CellTiter-Glo, of proliferating and senescent HT-1080$^{T}$ cells upon treatment with 5μM Navitoclax for 72 h. Senescence was induced with doxycycline (1 μg/ml) for 6 days. **(C)** Normalized viability, as evaluated by HTM-mediated quantification of nuclei, of proliferating and senescent HT-1080$^{T}$ cells upon treatment with GNF-7 at the indicated doses for 48 h. Senescence was induced with doxycycline (DOX: 1 μg/ml) or etoposide (ETO: 10 μM) for 6 days. **(D)** Time-point viability analysis of HT-1080$^{T}$ senescent cells after treatment with GNF-7 or Navitoclax at the indicated doses for 5 days. Senescence was induced with doxycycline (1 μg/ml) for 6 days.

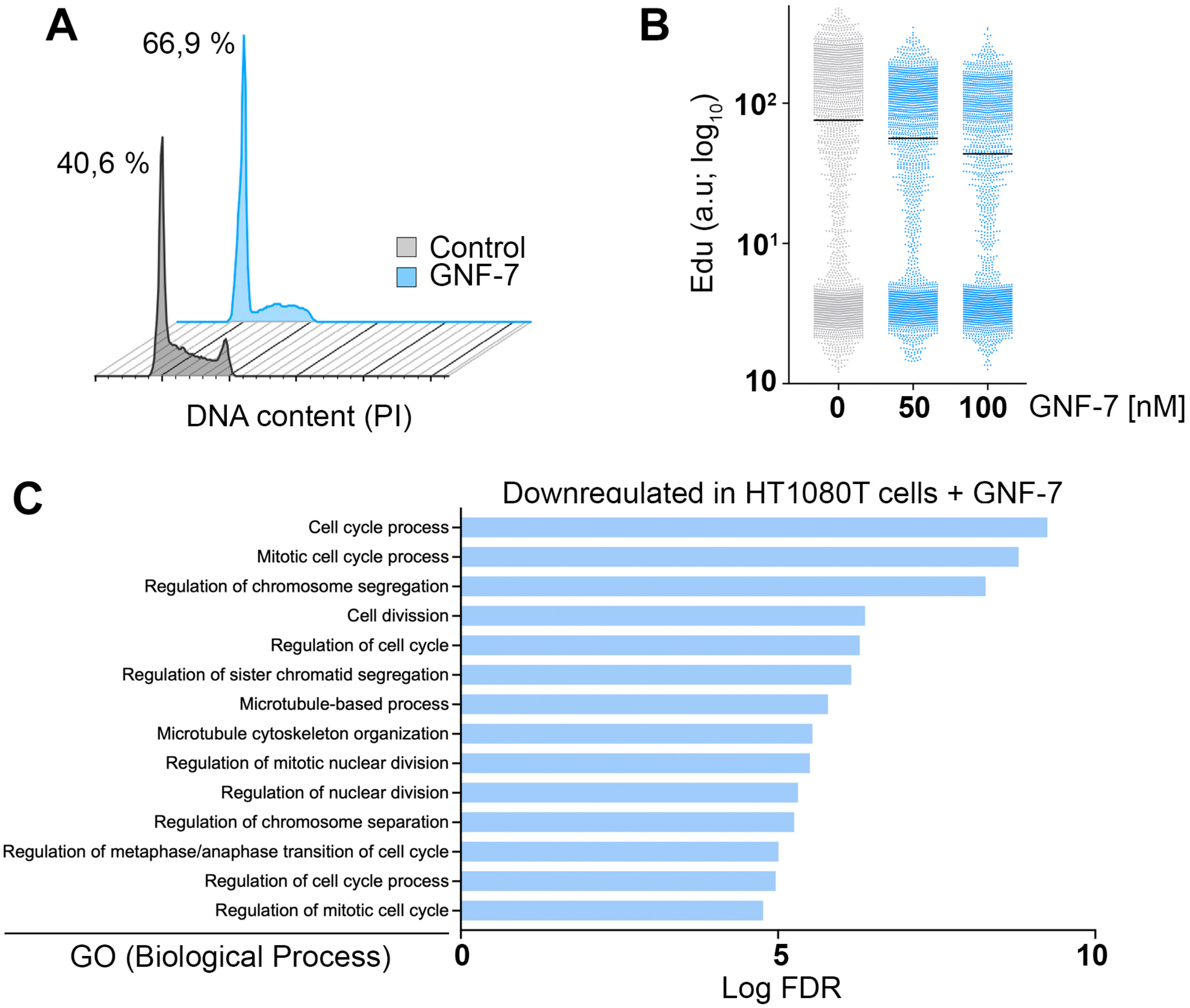
A
66,9 %
40,6 %
Control
GNF-7
DNA content (PI)
B
Edu (a.u; log10)
10²
10¹
10
0
50
100
GNF-7 [nM]
C
Downregulated in HT1080T cells + GNF-7
Cell cycle process
Mitotic cell cycle process
Regulation of chromosome segregation
Cell divission
Regulation of cell cycle
Regulation of sister chromatid segregation
Microtubule-based process
Microtubule cytoskeleton organization
Regulation of mitotic nuclear division
Regulation of nuclear division
Regulation of chromosome separation
Regulation of metaphase/anaphase transition of cell cycle
Regulation of cell cycle process
Regulation of mitotic cell cycle
GO (Biological Process)
0
5
10
Log FDR

**Figure S2. GNF-7 is cytostatic for proliferating cells.** (**A**) Cell cycle profile of proliferating HT-1080 cells untreated or treated with 50nM GNF-7 for 1 day. Numbers indicate the percentage of cells in G1. (**B**) EdU incorporation levels measured by HTM in proliferating HT-1080 cells treated with GNF-7 at the indicated doses. Black lines indicate median values. (**C**) GO analysis indicating the biological processes significantly associated with the downregulated genes in HT-1080 cells treated with 100nM GNF-7 for 8 h. The X-axis indicates $-\log_{10}$FDR, while Y-axis highlights the downregulated biological processes.

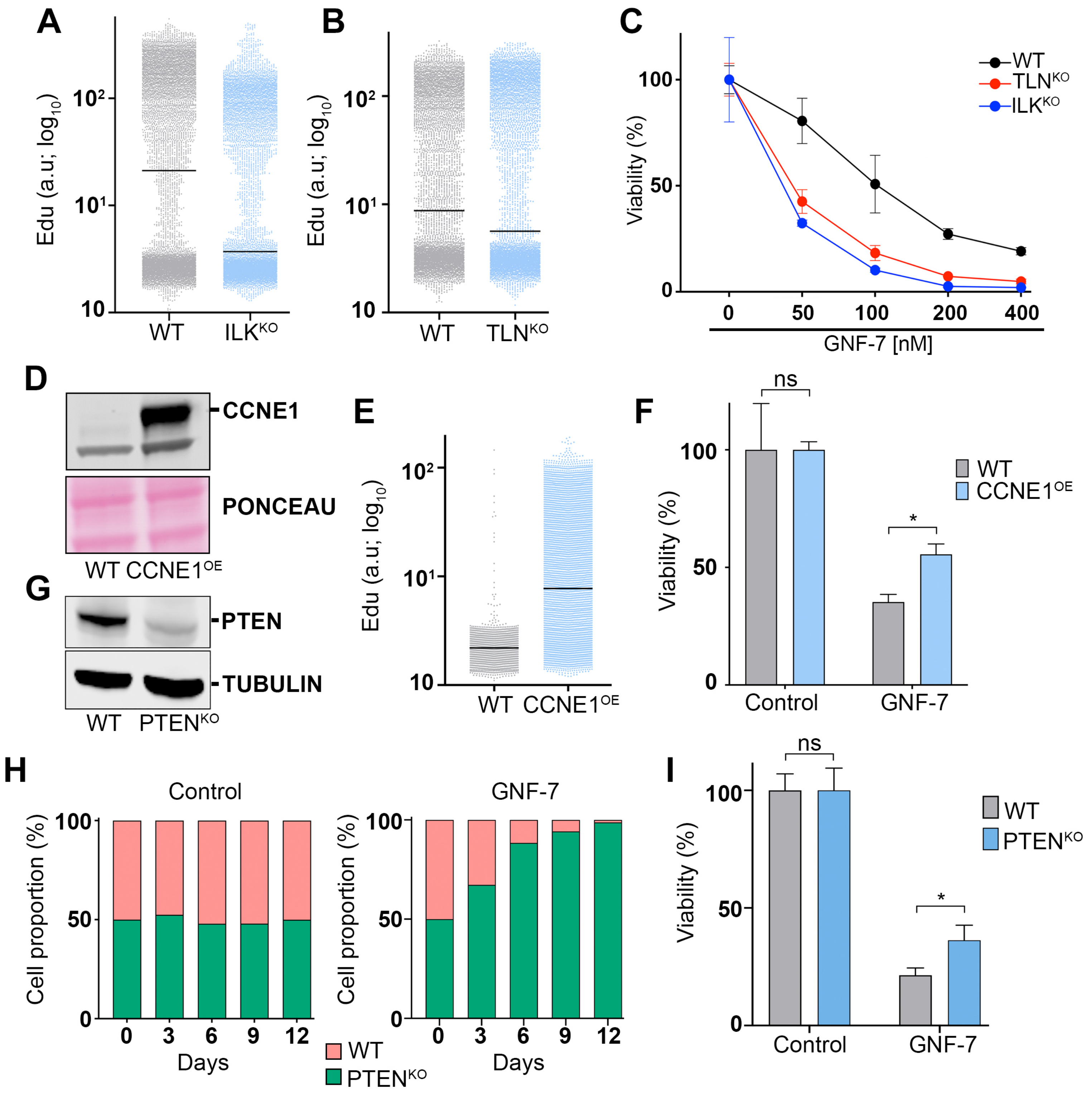
A
Edu (a.u; log10)
10
10¹
10²
WT
ILKKO
B
Edu (a.u; log10)
10
10¹
10²
WT
TLNKO
C
Viability (%)
0
50
100
0
50
100
200
400
GNF-7 [nM]
WT
TLNKO
ILKKO
D
CCNE1
PONCEAU
WT CCNE1OE
E
Edu (a.u; log10)
10
10¹
10²
WT
CCNE1OE
F
Viability (%)
0
50
100
ns
*
Control
GNF-7
WT
CCNE1OE
G
PTEN
TUBULIN
WT
PTENKO
H
Control
Cell proportion (%)
0
50
100
0
3
6
9
12
Days
GNF-7
Cell proportion (%)
0
50
100
0
3
6
9
12
Days
WT
PTENKO
I
Viability (%)
0
50
100
ns
*
Control
GNF-7
WT
PTENKO

**Figure S3. Mutations modulating the cellular response to GNF-7.** (**A, B**) EdU incorporation levels measured by HTM in proliferating WT, $ILK^{KO}$ and $TLN^{KO}$ HT-1080 cells. Black lines indicate median values. (**C**) Normalized viability of WT, $ILK^{KO}$ and $TLN^{KO}$ HT-1080 cells upon treatment with the indicated doses of GNF-7 for 48 h. (**D**) WB analysis illustrating the overexpression of CCNE1 present in $CCNE1^{OE}$ RPE cells. Ponceau levels are shown as a loading control. (**E**) EdU incorporation rates of WT and $CCNE1^{OE}$ RPE cells measured by HTM. Black lines indicate median values. (**F**) Normalized viability of WT and $CCNE1^{OE}$ RPE cells treated with 400nM GNF-7 for 48h, as measured by CellTiter-Glo. (**G**) WB of PTEN levels in WT and $PTEN^{KO}$ HT-1080 cells. TUBULIN levels are shown as a loading control. (**H**) Competition assay in wild type (red, WT) and PTEN-deficient (green, $PTEN^{KO}$) HT-1080 cells cultured in the presence or absence of 50 nM GNF-7. (**I**) Normalized viability, as measured by CellTiter-Glo, of senescent WT and $PTEN^{KO}$ HT-1080 cells after treatment with 200nM GNF-7 for 48 h. n.s.: non-significant, $^*P < 0.05$; *t*-test.

*Lopez et al* **FigS4**

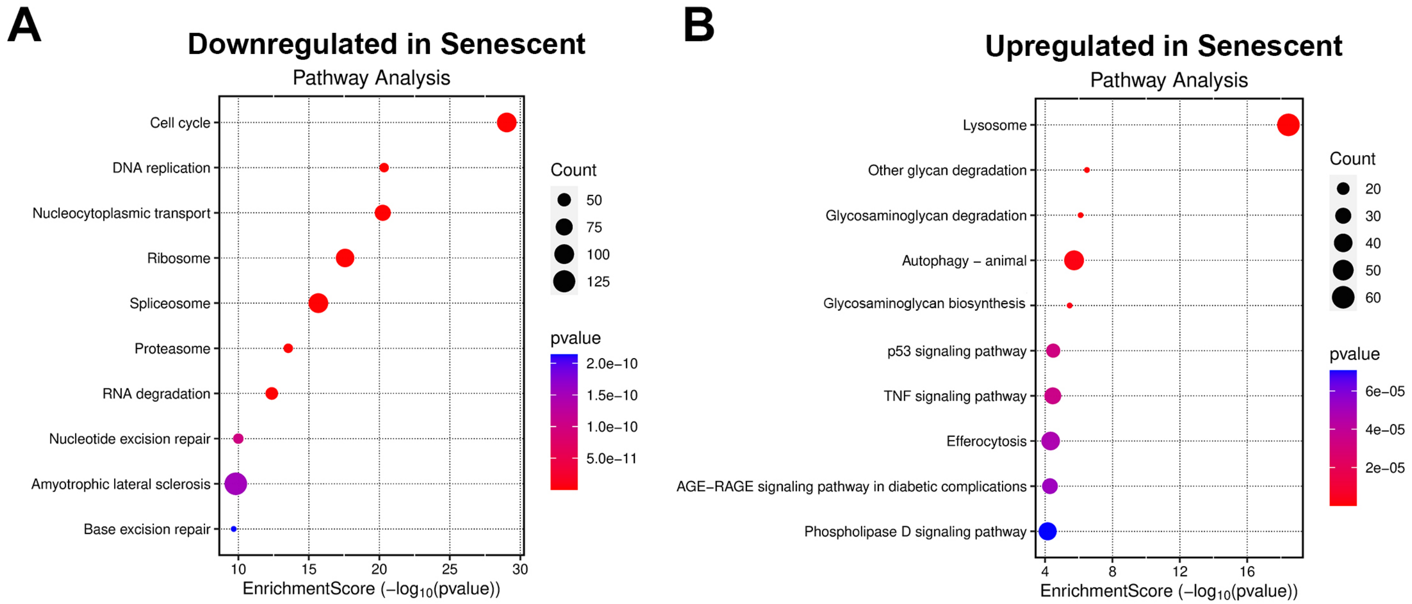

**Figure S4. Transcriptomic analyses in dox-treated HT-1080$^T$ cells.** (**A, B**) Dot plot representation of the significantly downregulated (**A**) or upregulated (**B**) pathways in senescent HT-1080$^T$ cells compared to proliferating ones. X-axis indicates the enrichment score (-Log10pvalue). Senescence was induced with doxycycline (1 μg/ml) for 6 days.

*Lopez et al* **FigS5**

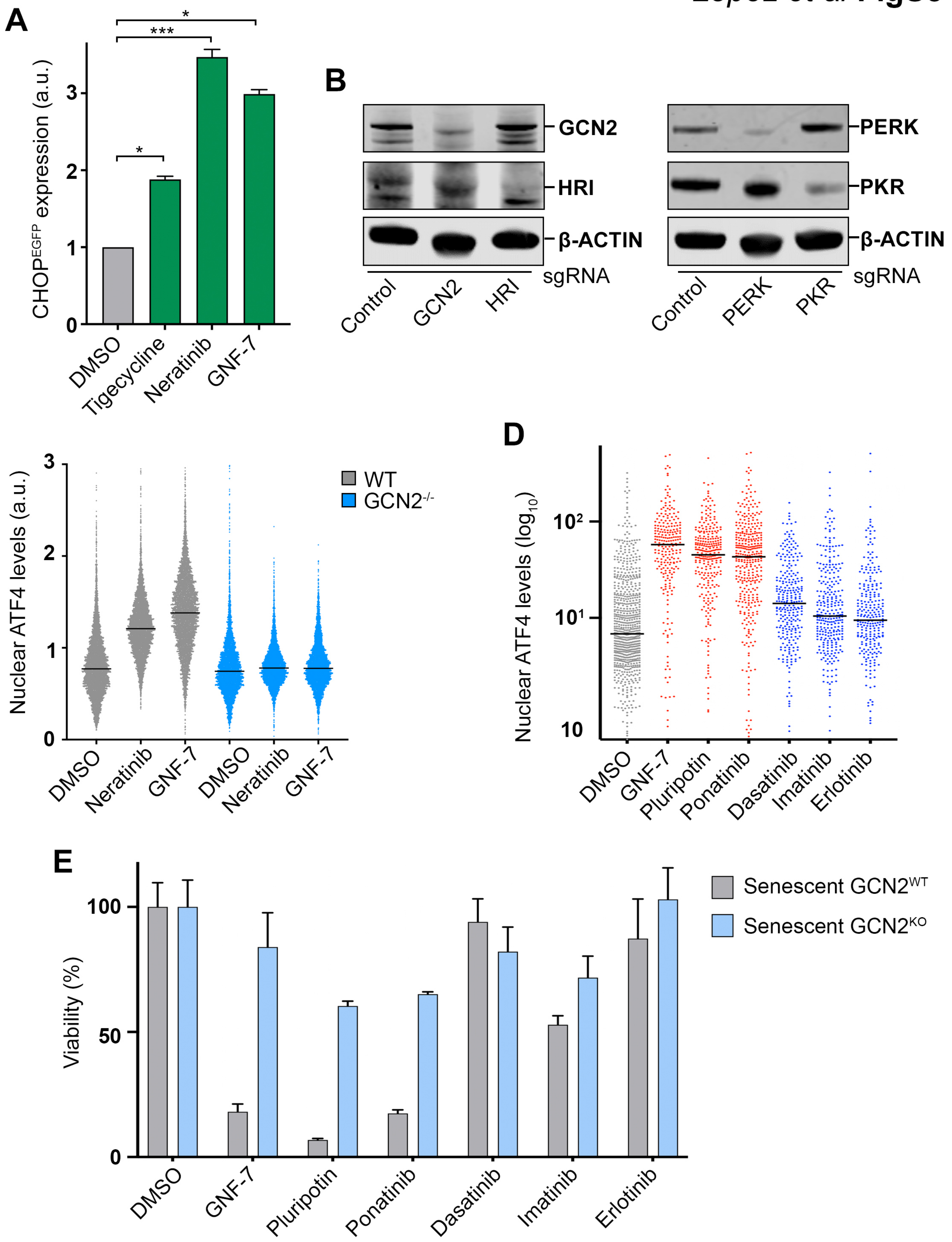

**Figure S5. GNF-7 and related TKis activate a GCN2-dependent ISR.** (**A**) Normalized EGFP levels, as quantified by flow cytometry, upon treatment of DLD1-$CHOP^{EGFP}$ cells with 10µM Tigecycline, 500nM Neratinib or 200nM GNF-7 for 72 h. (**B**) WB analyses of GCN2, HRI, PERK and PKR levels in DLD-1-$CHOP^{EGFP}$ cells where these genes were targeted by CRISPR-Cas9. β-ACTIN levels are shown as a loading control. (**C**) Nuclear ATF4 levels in WT and $GCN2^{KO}$ HT-1080 cells treated with the indicated compounds as measured by HTM. Cells were treated with 1µM Neratinib or 100nM GNF-7 for 16 h. Black lines indicate median values. (**D**) Nuclear ATF4 levels in senescent HT-1080 cells treated with the indicated drugs for 3 h. GNF-7 was added at 100nM, Erlotinib at 500nM, and the rest of drugs at 300nM. (**E**) Normalized viability, as measured by CellTiter-Glo, of senescent WT and $GCN2^{KO}$ HT-1080 cells treated with the indicated drugs for 48 h. The doses were the same as in (D). non-significant, *$P < 0.05$; ***$P < 0.001$; *t*-test.

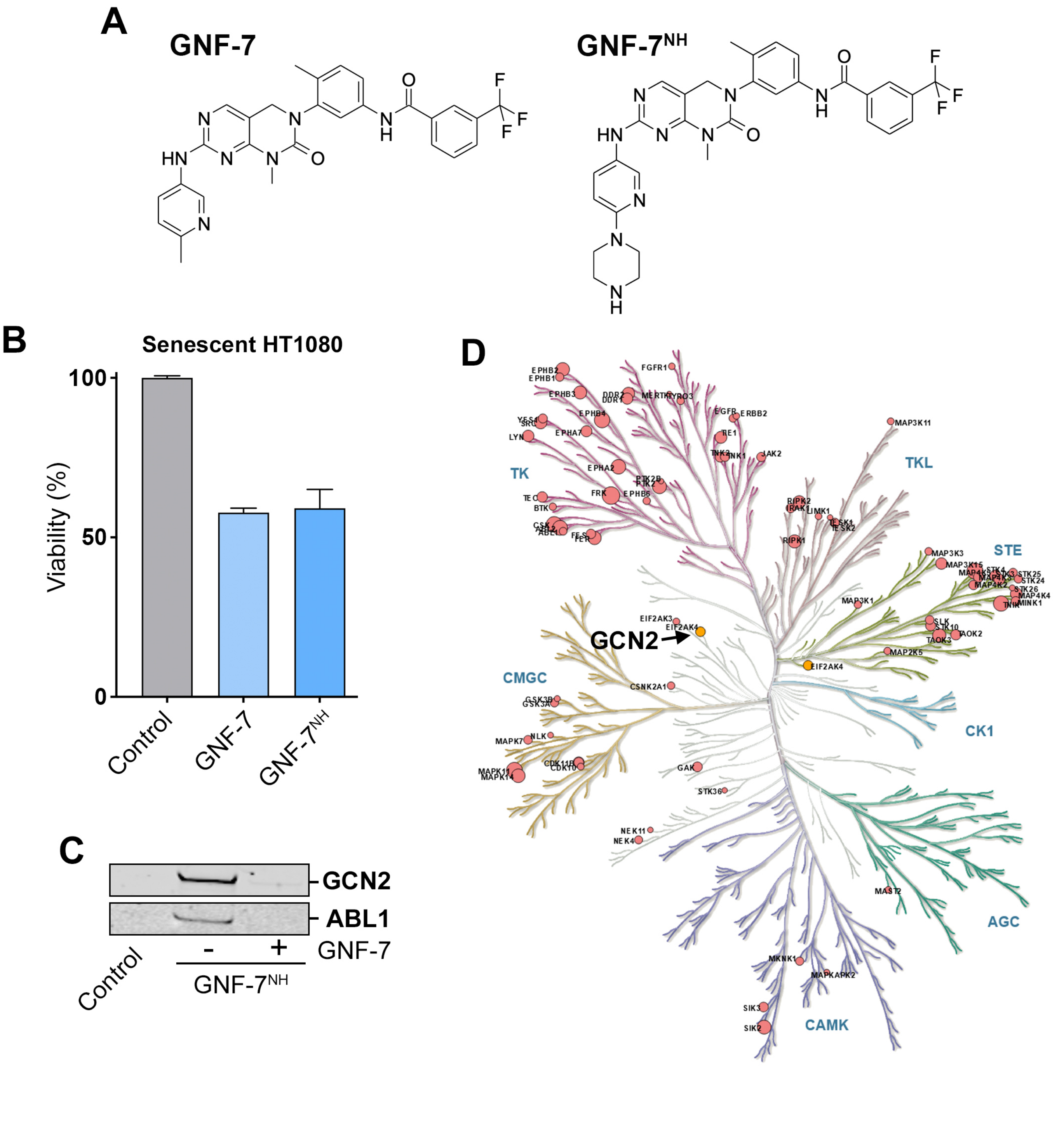
A
GNF-7
GNF-7NH
B
Senescent HT1080
Viability (%)
100
50
0
Control
GNF-7
GNF-7NH
C
GCN2
ABL1
Control
-
+
GNF-7
GNF-7NH
D
TK
TKL
STE
CK1
AGC
CAMK
CMGC
GCN2
EIF2AK4
EIF2AK3
EPHB2
EPHB1
EPHB3
EPHB4
EPHA7
EPHA2
FGFR1
DDR2
DDR1
TYRO3
EGFR
ERBB2
TIE1
TNK1
JAK2
FRK
EPHB6
BTK
LYN
SRC
MAP3K11
RIPK2
LIMK1
RIPK1
MAP3K3
MAP3K15
STK25
STK24
STK26
MAP4K4
MINK1
MAP4K2
MAP3K1
SLK
STK10
TAOK2
TAOK3
MAP2K5
CSNK2A1
GSK3B
GSK3A
NLK
MAPK7
MAPK14
GAK
STK36
NEK11
NEK4
MAST2
MKNK1
MAPKAPK2
SIK3
SIK2

**Figure S6. The landscape of GNF-7 interactors.** (**A**) Chemical structures of GNF-7 and its amine-derived analog, GNF-$7^{NH}$. (**B**) Normalized viability, as measured by CellTiter-Glo, of senescent HT-1080 cells treated with 400nM of GNF-7 or GNF-$7^{NH}$ for 72 h. Senescence was induced by palbociclib (5 μM) for 7 days. (**C**) WB analysis of the pull-down eluate of GNF-$7^{NH}$, illustrating the interaction of the drug with ABL1 and GCN2. The first lane (Control) represents uncoupled beads, while the second lane shows the interaction of GNF-$7^{NH}$ with its targets. The third lane presents the data upon competition with an excess of parental GNF-7 (100 μM, pre-treatment for 1 h). (**D**) Kinome map displaying the kinases identified as significantly interacting with GNF-$7^{NH}$. Circle sizes represent the log2FoldChange.

### GNF-7 derivatives

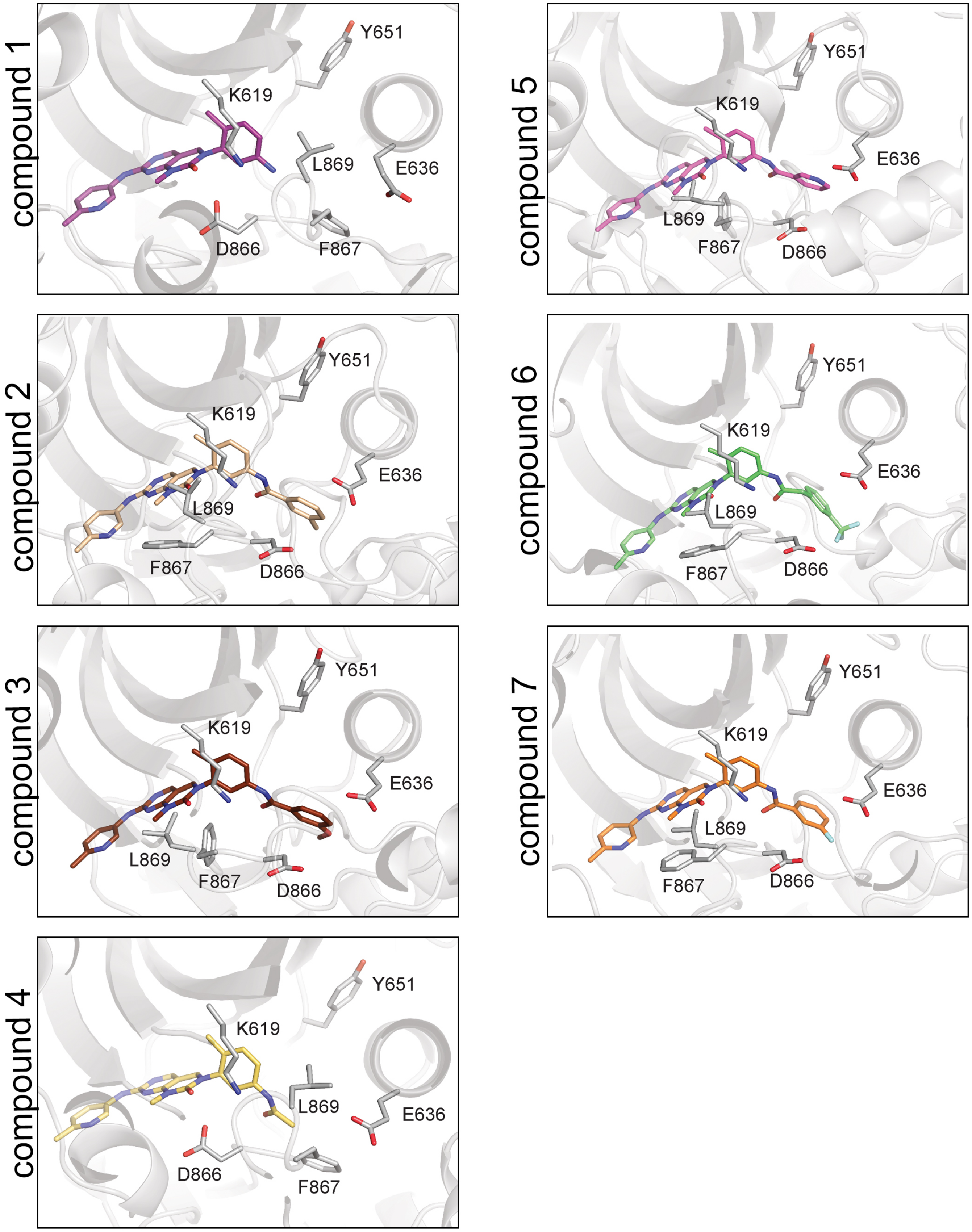

**Figure S7. Predicted structure of GNF-7 derivatives bound to GCN2.** Models of the binding of the GNF-7 derivatives shown in Fig. 6 to the kinase domain of GCN2. Docking analyses were performed using RoseTTAFold All-Atom (RFAA).

*Lopez et al* **FigS8**

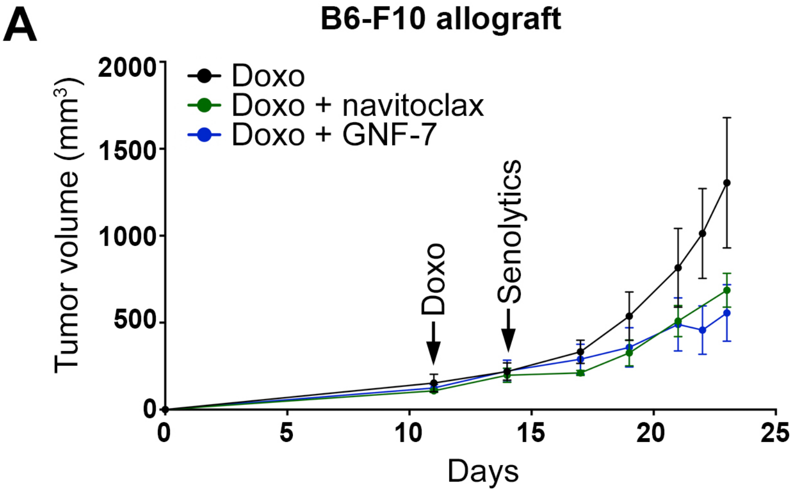


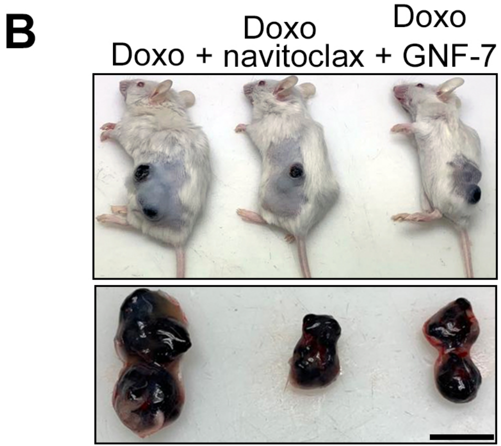

**Figure S8. GNF-7 synergizes with doxorubicin in a "one-two-punch" therapy of melanoma allografts.** (**A**) Tumour growth curves (mm3) of B16-F10 allografts upon treatment with doxorubicin alone, or doxorubicin followed by either navitoclax or GNF-7. Doxorubicin treatment started once the tumours reached an initial volume of around 150-200 mm3 and was added twice in alternate days to induce senescence (4mg/kg, intraperitoneal administration). After that, mice were treated with GNF-7 (15mg/kg, oral gavage) or Navitoclax (50mg/kg, oral gavage) every day. Arrows indicate the starting point of treatment: day 11 for doxorubicin and day 14 for Navitoclax and GNF-7 treatments. (**B**) Representative images of B16F10 allografts in mice at the endpoint of the tumour growth experiment (day 23). Scale bar (black) represents 1cm.

### Supplementary Tables and Videos

**Table S1**. Results from the primary chemical screen.

**Table S2**. Results from the dose-response validation screen.

**Table S3**. List of significant GNF-$7^{NH}$ interactors identified by proteomics.

**Table S4**. List of reagents used in this study.

**Videos S1, S2**

Time-lapse videos of senescent HT-1080 cells treated with DMSO as a control (**S1**) or 200nM of GNF-7 (**S2**) for 48h. Apoptosis was monitored with a fluorescent reporter that detects the activation of caspase 3 (green). Nuclei were visualized with Hoechst 33342 (blue).

**Videos S3, S4**

Time-lapse videos of senescent wild type (**S3**) or GCN2-deficient (**S4**) HT-1080 cells treated with 200nM of GNF-7 for 48h. Apoptosis was monitored with a fluorescent reporter that detects the activation of caspase 3 (green). Nuclei were visualized with Hoechst 33342 (blue).
