## Supplementary material for "The tyrosine kinase inhibitor GNF-7 targets senescent cells through allosteric activation of GCN2": Table S4

**Table S4. List of reagents used in this study**

| **REAGENT or RESOURCE** | **SOURCE** | **IDENTIFIER** |
| --- | --- | --- |
| Antibodies | | |
| Rabbit anti-ATF4 | Cell Signalling | 11815 |
| Rabbit anti-eIF2α | Cell Signaling | 9722 |
| Rabbit anti-eIF2α-phospho | Thermo Fisher | 44-728G |
| Mouse anti-CHOP | Cell Signalling | 3302 |
| Rabbit anti-GCN2 | Cell Signalling | 3302 |
| Rabbit anti-GCN2-phospho (T899) | Abcam | 75836 |
| Rabbit anti-HRI | MyBioSource | MBS2538144 |
| Mouse anti-PERK | Santa Cruz | SC-377400 |
| Mouse anti-PKR | Santa Cruz | SC-6282 |
| Mouse anti-actin | Sigma | A5441 |
| Mouse anti-Cas9 | Cell Signalling | 14697 |
| Mouse anti -Tubulin | Sigma | T9026 |
| Rabbit ant- FLAG | Sigma | F1804 |
| Mouse anti-PTEN | Santa Cruz | sc-7974 |
| Mouse anti-TSC1 | Santa Cruz | sc-377386 |
| Rabbit anti-p21 | Cell Signaling | 2947 |
| Mouse anti-CCNE1 | Santa Cruz | sc-247 |
| Mouse anti-TLN1 | Santa Cruz | sc-365875 |
| Rabbit anti-NF2 | Proteintech | 21686-1-AP |
| Mouse anti-ABL1 | Proteintech | 68254-1-Ig |
| Rabbit anti-p38-phospho | Cell Signaling | 4631 |
| Anti-Mouse IgG-488 | Invitrogen | Cat#A11001 |
| Anti-Mouse IgG-555 | Bethyl | Cat#A90-516D3 |
| Anti-Mouse IgG-647 | Invitrogen | Cat#A21463 |
| Anti-Rabbit IgG-488 | Invitrogen | Cat#A21441 |
| Anti- Rabbit IgG-555 | Bethyl | Cat#A120-201D3 |
| Anti- Rabbit IgG-647 | Invitrogen | Cat#A21443 |
| Chemicals, peptides, and recombinant proteins |  |  |
| CellTiter-Glo | Promega | G7571 |
| 4-Hydroxytamoxifen (4 OHT) | Sigma | H7904 |
| Doxorubicin | Sigma | D1515 |
| Doxycycline | Pancreac Applichem | D9891 |
| Erlotinib | Selleckchem | S7786 |
| Navitoclax | MedChemExpress | HY-10087 |
| Pan Caspase Inhibitor Z-VAD-FMK | Bio-techne R&D SYSTEMS S.L | FMK001 |
| GNF-7 | MedChemExpress | HY-10943 |
| Pluripotin | MedchemExpress | HY-10579 |
| Ponatinib | MedChemExpress | HY-12047 |
| Dasatinib | MedChem Express | HY-10181 |
| Imatinib | MedChem Express | HY-15463 |
| Palbociclib | MedChem Express | HY-50767 |
| ABT-263 (Navitoclax) | MedChem Express | HY-10087 |
| Capivasertib | MedChem Express | HY-15431 |
| Hydroxyurea (HU) | Sigma | H8627 |
| Nocodazole | Sigma | M1404 |
| Thymidine | Sigma | T-1895 |
| ISRIB | Sigma | SML0843 |
| Puromycin | Sigma | P8833 |
| Blasticidin | Life Technologies | A11139-03 |
| Tigecycline | Sigma | Y0001961 |
| L-mimosine | Sigma | M0253 |
| Primocin | Invivogen | ant-pm-1 |
| N-Acetylcysteine | Sigma | A9165 |
| Nicotinamide | Sigma | N0636 |
| Noggin | Gibco | 17811063 |
| Neuregulin | Peprotech | 15144093 |
| EGF | StemCell Tech. | 78006.1 |
| FGF-7 | Peprotech | 100-19 |
| FGF-10 | Peprotech | 100-26 |
| B27 Suplement | Gibco | 17504044 |
| R-Spondin 3 | Gibco | 120-44 |
| A83-01 | Tocris | 2939 |
| SB202190 | Sigma | S7067 |
| Y-27632 | Axon Medchem | 129830-38-2 |
| Triethylamine | Sigma, | T0886 |
| Ethanolamine | Sigma | 11016-7 |
| Critical commercial assays |  |  |
| Click-It EU 488 Imaging kit | Thermo Fisher scientific | C10329 |
| Click-iT EdU Cell Proliferation Assay kit | Thermo Fisher scientific | C10337 |
| Absolutely RNA Microprep kit | Agilent | 400805 |
| Gentra Puregene Blood Kit | Qiagen | 158445 |
| KAPA HIFI Hot Start PCR kit | Roche | KK2502 |
| RNA Clean & Concentrator-5 kit | Zymo Research | R1013 |
| Lipofectamine 2000 | Invitrogen | Cat#11668027 |
| QuantSeq 3' mRNA-Seq Library Prep Kit | Lexogen | N/A |
| Lipofectamine RNAiMAX | Thermo Fisher Scientific | 13778150 |
| senescence-associated β-galactosidase kit | Cell Signaling | 9860 |
| NEB® Golden Gate Assembly Kit (BsmBI-v2) | NEB | 174E1602S |
| APC Annexin V | BD Biosciences | 550474 |
| NHS-Activated Sepharose 4 Fast Flow Beads | GE LifeSciences | 17090601 |
| Mini Bio-Spin Chromatography Columns | BioRad | 7326207 |
| Deposited data |  |  |
| Proteomic Data | This paper |  |
| RNA-seq data | This paper |  |
| Experimental models: Cell lines |  |  |
| HT1080 | ATCC | N/A |
| MCF7 | ATCC | N/A |
| DLD-1 | ATCC | N/A |
| RPE | ATCC | N/A |
| B16-F10 | ATCC | N/A |
| KBM7 | ATCC | N/A |
| mESCs | ATCC | N/A |
| RPE hTERT^Cas9^ TP53^-/-^ CCNE1^OE^ | Dr. Daniel Durocher’s laboratory |  |
| Patient-derived ER+ breast cancer organoids | Dr. Miguel Quintela’s laboratory | N/A |
| Experimental models: Organisms/strains |  |  |
| Mouse: C57BL/6 mice | This paper | N/A |
| severe combined immunodeficient (Prkdc^scid^) | This paper | N/A |
| Oligonucleotides. sgRNAs for CRISPR/Cas9 gene editing: |  |  |
| PTEN: TCATCTGGATTATAGACCAG | This paper | N/A |
| NF2: CATCTCGTACAGTGACAAGG | This paper | N/A |
| ILK_CCCCTGAGAGAGCTTCTCCG | This paper | N/A |
| GCN2: TCGTTGCGGGTAGCTCTC | This paper | N/A |
| TSC1: TGTTTACAAGCATAGGGCCA | This paper | N/A |
| USP48: AGACTGTGGATCTTTCACGT | This paper | N/A |
| CCNC: ATTTACGTGAACACACAGCA | This paper | N/A |
| TLN1: GGATCCGCTCACGAATGATG | This paper | N/A |
| BCL2: AGGAGAAGATGCCCGGTGCG | This paper | N/A |
| GCN1: AAAGAAATCCTATACCTGAG | This paper | N/A |
| HRI: GCGGGAAAGTCGATGGCCGG | This paper | N/A |
| PERK: CTCAGCGACGCGAGTACCGG | This paper | N/A |
| PKR AATACATACCGTCAGAAGCA | This paper | N/A |
| PTK2: ATCAGTTACCTAACGGACAA | This paper | N/A |
| Recombinant DNA |  |  |
| plentiCRISPR v2-Blast | Addgene | (Plasmid #83480) |
| plentiCRISPRv2-puro | Addgene | (Plasmid #52961) |
| pLentiGuide-mCherry | Gift by Dr Cristina Mayor-Ruiz | N/A |
| pLentiGuide-GFP | Gift by Dr Cristina Mayor-Ruiz | N/A |
| plentiCas9-Blast | Gift by Dr Cristina Mayor-Ruiz | N/A |
| pCLX‐CHOP‐dGFP | Addgene | (Plasmid #71299) |
| Mouse Toronto Knockout (mTKO) CRISPR Library | Addgene | (Plasmid #159393, two-vector system) |
| Software and algorithms |  |  |
| CRISPOR tool |  | <https://crispor.gi.ucsc.edu/crispor.py> |
| inDelphi | Shen et al., *Nature*, 2019 | <https://indelphi.giffordlab.mit.edu/> |
| Gene Ontology | <https://geneontology.org/> |  |
| GraphPad Prism 9 | GraphPad Software Inc | http://www.graphpad.com/scientific-software/prism/ |
| Model-based Analysis of Genome-wide CRISPR/Cas9 Knockout (MAGeCK) | Li et al., *Genome Biol*., 2014 |  |
| MaGIC Volcano Plot Tool |  | https://volcano.bioinformagic.tools |
| RoseTTAFold All-Atom (RFAA) | Marchal I., *Nature Biotechnology*, 2024 |  |
| PyMOL 3.1 |  | Schrödinger et al., *PyMOL,* 2020 |
| Gene Expression Profiling Interactive Analysis: GEPIA2 |  | <http://gepia2.cancer-pku.cn/> |

**Patient-derived organoid media: Ad-DF+++ medium.**

| **Factor** | **Supplier** | **Catalogue nº** | **Final concentration** |
| --- | --- | --- | --- |
| Advanced DMEM/F12 | Gibco | 12634010 | 1X |
| HEPES | Thermo Scientific™ | 35050038 | 10 mM |
| GlutaMax | Gibco | 15630049 | 1X |
| Penicillin/Streptomycin | Gibco, | 11548876 | 100 U/ml |
| Primocin | Invivogen | ant-pm-1 | 50 µg/ml |

**Patient-derived organoid media: Complete organoid medium.**

| **Factor** | **Supplier** | **Catalogue nº** | **Final concentration** |
| --- | --- | --- | --- |
| Advanced DMEM/F12 | Gibco | 12634010 | 1X |
| Hepes | Thermo Scientific™ | 35050038 | 10 mM |
| GlutaMax | Gibco | 15630049 | 1X |
| Penicillin/Streptomycin | Gibco, | 11548876 | 100 U/ml |
| Primocin | Invivogen | ant-pm-1 | 50 µg/ml |
| N-Acetylcysteine | Sigma | A9165 | 1.25 mM |
| Nicotinamide | Sigma | N0636 | 5 mM |
| Noggin | Gibco | 17811063 | 100 ng/ml |
| Neuregulin | Peprotech | 15144093 | 5 nM |
| EGF | StemCell Tech. | 78006.1 | 2,5 ng/ml |
| FGF-7 | Peprotech | 100-19 | 5 ng/ml |
| FGF-10 | Peprotech | 100-26 | 20 ng/ml |
| B27 Suplement | Gibco | 17504044 | 1X |
| R-Spondin 3 | Gibco | 120-44 | 63 ng/ml |
| A83-01 | Tocris | 2939 | 500nM |
| SB202190 | Sigma | S7067 | 500 nM |
| Y-27632 | Axon Medchem | 129830-38-2 | 5µM |
